# Microenvironmental Sensing by Fibroblasts Controls Macrophage Population Size

**DOI:** 10.1101/2022.01.18.476683

**Authors:** Xu Zhou, Ruth A. Franklin, Miri Adler, Trevor S. Carter, Emily Condiff, Taylor S. Adams, Scott D. Pope, Naomi H. Philip, Matthew L. Meizlish, Naftali Kaminski, Ruslan Medzhitov

**Affiliations:** Department of Immunobiology, Yale University School of Medicine, New Haven, Connecticut 06510, USA; Division of Gastroenterology, Hepatology and Nutrition, Department of Pediatrics, Boston Children’s Hospital and Harvard Medical School, Boston, Massachusetts 02115, USA; Department of Stem Cell and Regenerative Biology, Harvard University, Cambridge, Massachusetts 02138, USA; Department of Immunology, Harvard Medical School, Boston, Massachusetts 02115, USA; Broad Institute of Massachusetts Institute of Technology and Harvard, Cambridge, Massachusetts 02142, USA; Department of Electrical Engineering and Computer Science, Massachusetts Institute of Technology, 77 Massachusetts Ave, Cambridge mA 02139; Pulmonary Critical Care and Sleep Medicine, Yale University School of Medicine, New Haven, Connecticut 06510, USA; Howard Hughes Medical Institute, Yale University School of Medicine, New Haven, Connecticut 06510, USA; Department of Medicine, Massachusetts General Hospital, Boston, MA, 02114, USA

**Keywords:** Cell composition, CSF1, fibroblasts, macrophages, YAP1, SMAD

## Abstract

Animal tissues are comprised of diverse cell types. However, the mechanisms controlling the number of each cell type within tissue compartments remain poorly understood. Here, we report that different cell types utilize distinct strategies to control population numbers. Proliferation of fibroblasts, stromal cells important for tissue integrity, is limited by space availability. In contrast, proliferation of macrophages, innate immune cells involved in defense, repair, and homeostasis, is constrained by growth factor availability. Examination of density-dependent gene expression in fibroblasts revealed that Hippo and TGF-*β* target genes are both regulated by cell density. We found YAP1, the transcriptional co-activator of the Hippo signaling pathway, directly regulates expression of *Csf1*, the lineage-specific growth factor for macrophages, through an enhancer of *Csf1* that is specifically active in fibroblasts. Activation of YAP1 in fibroblasts elevates *Csf1* expression and is sufficient to increase the number of macrophages at steady state. Our data also suggest that expression programs in fibroblasts that change with density may result from sensing of mechanical force through actin-dependent mechanisms. Altogether, we demonstrate that two different modes of population control are connected and coordinated to regulate cell numbers of distinct cell types. Sensing of the tissue environment may serve as a general strategy to control tissue composition.

**Significance Statement:** Collections of distinct cell types constitute animal tissues. To perform their unique functions, each cell type must exist in the correct number and proportion in a given tissue compartment. However, many of the mechanisms regulating and coordinating cell population sizes remain enigmatic. Our study characterizes two different modes of population size control, utilized by two ubiquitous cell types, macrophages and fibroblasts. Macrophage populations are more sensitive to the presence of growth factors in the environment and fibroblasts are more sensitive to space limitations. Intriguingly, space-sensing mechanisms in fibroblasts directly control the production of growth factor for macrophages and thus macrophage numbers. This link suggests a mechanism by which macrophage compartment size is controlled by stromal cells according to the microenvironment.

## Introduction

Animal tissues consist of multiple cell types present in appropriate numbers and ratios (Raff, 1996). Each cell type requires a specific growth factor for survival and proliferation, and therefore, local availability of growth factors can control cell numbers within tissue compartments (Raff, 1992). However, growth factors alone are often insufficient to determine tissue compartment size (Hart et al., 2012). In fact, the maximum population size that can be supported in a particular environment (the carrying capacity of the environment), is determined by any limiting factor of that environment, such as space, nutrients, and oxygen levels (Gotelli, 2008). The interplay between growth factor availability and the carrying capacity of tissue environment in defining compartment size is not well understood. In particular, it is not well known how the numbers of different cell types within tissue compartments are maintained and coordinated (Meizlish et al., 2021). Factors that control compartment size, and ultimately tissue and organ size, generally fall into two categories (Figure 1A): First, compartment size can be controlled by the tissue micro-environment (Duronio and Xiong, 2013). One of the best understood examples is space availability, which is sensed through mechanical properties of the environment, including physical contact with extracellular matrix (ECM) or neighboring cells (Chen et al., 1997; Humphrey et al., 2014; Hynes and Naba, 2012; Nelson et al., 2005). In this scenario, cells proliferate until available space is used up, at which point, cell-ECM or cell-cell contacts suppress further cell division to reach a maximal cell number in a given environment. This space-dependent constraint can be sensed by signaling pathways, such as the Hippo signaling pathway (Dupont et al., 2011; Gumbiner and Kim, 2014; Yu et al., 2015) and mechanosensitive GPCRs (Gudipaty et al., 2017), and is reflected in *in vitro* phenomenon of ‘contact inhibition’ of growth (Eagle and Levine, 1967). However, mechanosensing of space availability is not a usable strategy for some cell types, including hematopoietic cells, which are not constrained by space. Thus, a second strategy to control compartment size is through the availability of lineage-specific growth factors (Raff, 1992). Indeed, the numbers of naïve and memory T cells is maintained by IL-7 (Bradley et al., 2005; Schluns et al., 2000) and the number of macrophages is limited by CSF1 (Chitu and Stanley, 2006; Guilliams et al., 2020; Zhou et al., 2018). In these examples, the cell numbers depend on the local availability of appropriate growth factors and therefore, population size in a tissue compartment is limited by growth factor availability, rather than space availability (Buechler et al., 2021). According to this paradigm, if cell numbers are above or below the level that can be supported by available growth factors, cells will either die or proliferate, respectively, until they reach the steady state (Raff, 1992).

**Figure 1.**
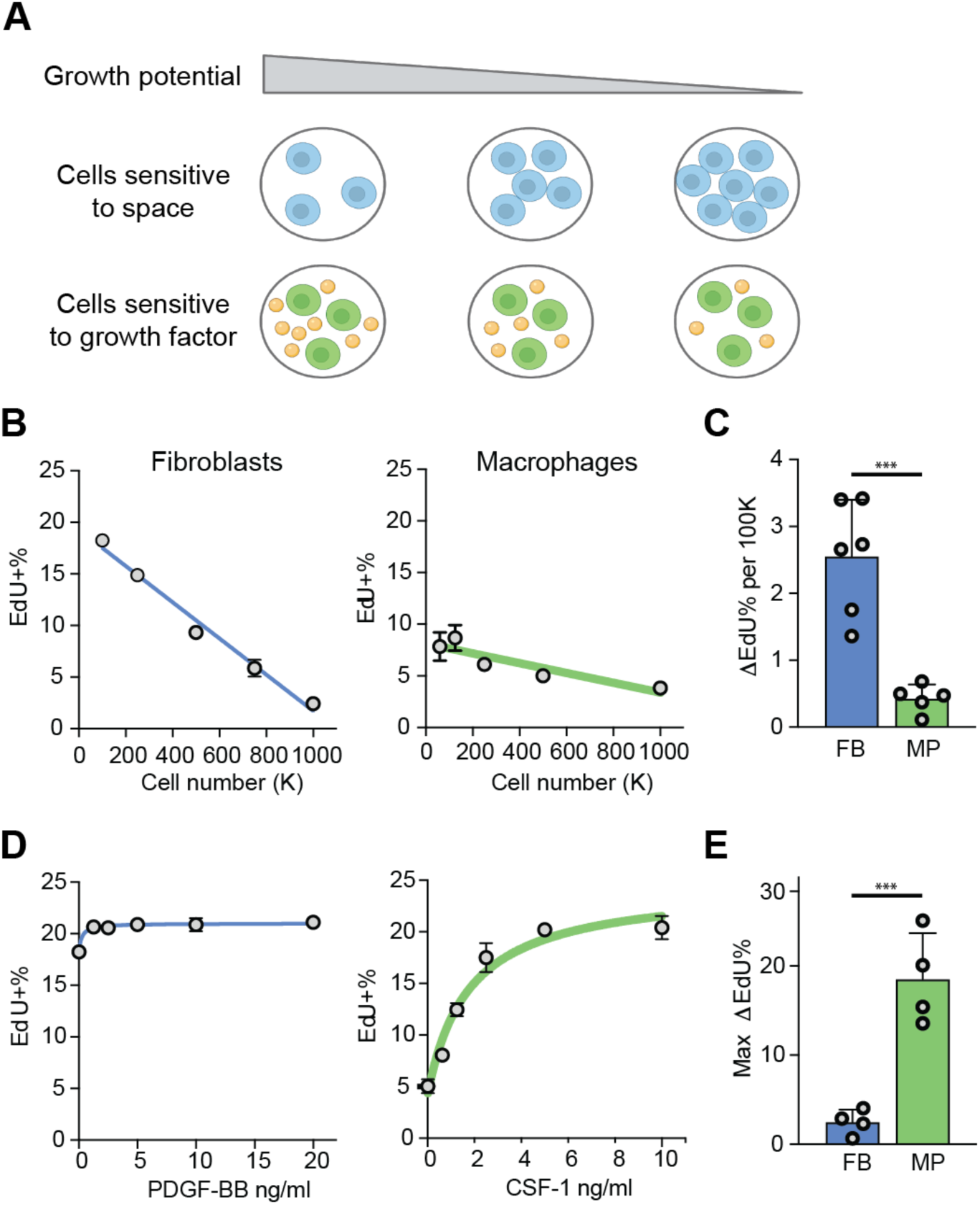
Fibroblasts and macrophages use two distinct mechanisms to control cell numbers. A. Two proposed mechanisms of cell number control in a given tissue compartment. Orange spheres represent growth factors. B. Proliferation of BMDMs and MEFs, measured as the percentage of EdU^+^ cells after 2 hr of EdU labeling, following overnight culture at the indicated densities. C. Cell density-dependent proliferation, estimated based on the linear fit of density-dependent growth as in (B) and quantified as the differential proliferation per 100,000 cells. Each dot represents an independent experiment. D. Proliferation of BMDMs and MEFs stimulated with recombinant CSF-1 or PDGF-BB respectively, measured as the percentage of EdU^+^ cells after 2 hr of EdU labeling, cultured at the indicated growth factor concentration overnight. E. Growth factor dependent proliferation, quantified as the maximum change of EdU^+^ cells with the addition of growth factors.

Although for a given cell type either space or growth factor availability can be the dominant factor controlling cell numbers, any factor required for cell proliferation can impose a limit on population size. Thus, even growth of cell types normally restricted by space availability can become dependent on growth factor availability if the growth factor rather than space becomes a limiting factor. Indeed, some cell types can be limited by either space of growth factor availability depending on circumstances. For instance, proliferation of hepatocytes is inhibited by Hippo-YAP signaling, (Lu et al., 2010; Su et al., 2015; Zhang et al., 2010) while also regulated by hepatic growth factors (Nakamura et al., 2011). Intestinal epithelial cells are sensitive to mechanical forces sensed by PIEZO1 (Gudipaty et al., 2017) and are also dependent on several growth factors such as EGF and WNTs (Sato et al., 2011). Furthermore, different cell types within the same tissue compartment may employ different strategies to control each of their population numbers. It is unclear how cells using these two strategies coordinate their numbers within tissues. Since tissues are made up of multiple cell types, it is unknown whether sensing of environmental cues in one cell type would influence the population size of another cell type. Addressing these questions will be important for understanding of how different types of cells collectively constitute an organized and functional tissue.

Within tissues, growth factor for one cell type is typically produced by another cell type as a paracrine signal. Previously, we found that macrophages and fibroblasts, two cell types present in most mammalian tissues (Guilliams et al., 2020; Lynch and Watt, 2018), exchange growth factor signals PDGFB and CSF1 *in vitro* (Zhou et al., 2018). This reciprocal communication through growth factors supports stable population ratios of these two cell types (Adler et al., 2018; Buechler et al., 2021; Franklin, 2021; Zhou et al., 2018). Similar paracrine production of CSF1 by fibroblasts for macrophages has been demonstrated *in vivo* in liver (Bellomo et al., 2020) and spleen (Bonnardel et al., 2019). Here, we examined how different cell number control mechanisms apply to these two cell types, and found that at steady state, fibroblast proliferation is constrained by space availability, while macrophage proliferation is dependent on growth factor availability. Moreover, we found that sensing space availability in fibroblasts through YAP1 directly controls expression of the macrophage-specific growth factor CSF1. YAP1 activity in fibroblasts controls the absolute number of macrophages, as well as the ratios between these two cell types. We propose that the two strategies of cell number control, by space and growth factor availability, are inherently linked through the regulation of paracrine growth factor signals. This mechanism may provide a simple solution for automatically adjusting cell numbers and ratios within tissues.

## Results

### Fibroblasts and macrophages employ different modes of compartment size control

We previously observed that proliferation of fibroblasts, but not macrophages, is constrained by space availability: even in the presence of growth signals, space constraints set the carrying capacity of fibroblasts *in vitro* (Zhou et al., 2018). We thus hypothesized that fibroblasts and macrophages may control cell numbers using different strategies. First, we formally examined whether fibroblast proliferation is more sensitive to space constraint, by measuring proliferation rates of bone marrow-derived macrophages (BMDMs, prototypical macrophages) and mouse embryonic fibroblasts (MEFs, prototypical fibroblasts) at different cell densities. The thymidine analog EdU is incorporated into newly synthesized DNA during S phase, allowing proliferating cells to be identified. The percentage of EdU^+^ cells in a short time window (2 hours) was thus used as a proxy for the proliferation rate. Proliferation of fibroblasts was more density-dependent compared to proliferation of macrophages (Figure 1B & 1C). Second, we tested how growth rates of macrophages and fibroblasts depend on growth factor availability. We titrated the amounts of CSF1 for macrophages and PDGFB for fibroblasts and observed that addition of growth factors had a bigger impact on proliferation of macrophages compared to fibroblasts (Figure 1D & 1E). Interestingly, even though proliferation of fibroblasts requires growth factors, in all conditions we have examined, space constraints regulate proliferation of fibroblasts almost five times more than the addition of growth factors; in contrast, growth factors influence proliferation of macrophages about five times more than changes in cell density. These data demonstrate two modes of compartment size control used by different cell types: responsiveness to environmental limitations such as space availability, as observed for fibroblasts, and responsiveness to growth factor availability, as observed for macrophages.

These findings raise a question: If different cell types employ these distinct strategies to control their cell numbers, how is sensing of the environment and production of growth factors coordinated to achieve appropriate cellular composition of tissues? In a previous study, we found that macrophages and fibroblasts interact in a stable circuit via cell-cell contact and production of growth factors (Zhou et al., 2018). While growth factors are required to maintain both populations, the stable number of fibroblasts is determined by space as the limiting factor. We thus hypothesized that these two distinct modes of population control do not operate independently, but rather are coupled and coordinated, to control cell numbers within tissues. Specifically, we hypothesized that sensing of space availability by fibroblasts is translated into growth factor production for macrophages, thereby adjusting cell numbers controlled by these two mechanisms.

### Fibroblasts display density-dependent gene expression programs

To test this hypothesis, we first evaluated cell density-dependent gene expression in fibroblasts. RNAseq was performed on fibroblasts at four cell densities, using two batches of separately isolated primary fibroblasts. The highest cell density is close to the theoretical carrying capacity estimated previously (highly confluent) (Zhou et al., 2018), and the lowest cell density is close to the threshold at which fibroblasts cannot survive in monoculture (sparse). At each cell density, gene expression of biological replicates is highly correlated (Figure S1A). To select bona fide density-dependent genes, we determined differentially expressed genes, taking into account biological variation, and developed an automated algorithm to identify the genes with consistent expression changes across two data sets (see Methods, Figure S1B, C). In total, we found 1,826 genes to be highly expressed (TPM > 2) and significantly regulated by cell density in fibroblasts. Using unsupervised K-mean clustering, these genes are grouped into seven groups with distinct patterns of gene expression (Figure 2A). Three clusters are up-regulated at low cell density (L1-L3), and four clusters are up-regulated at high cell density (H1-H4). L1 and H1 contain genes that are the most differentially regulated.

**Figure 2.**
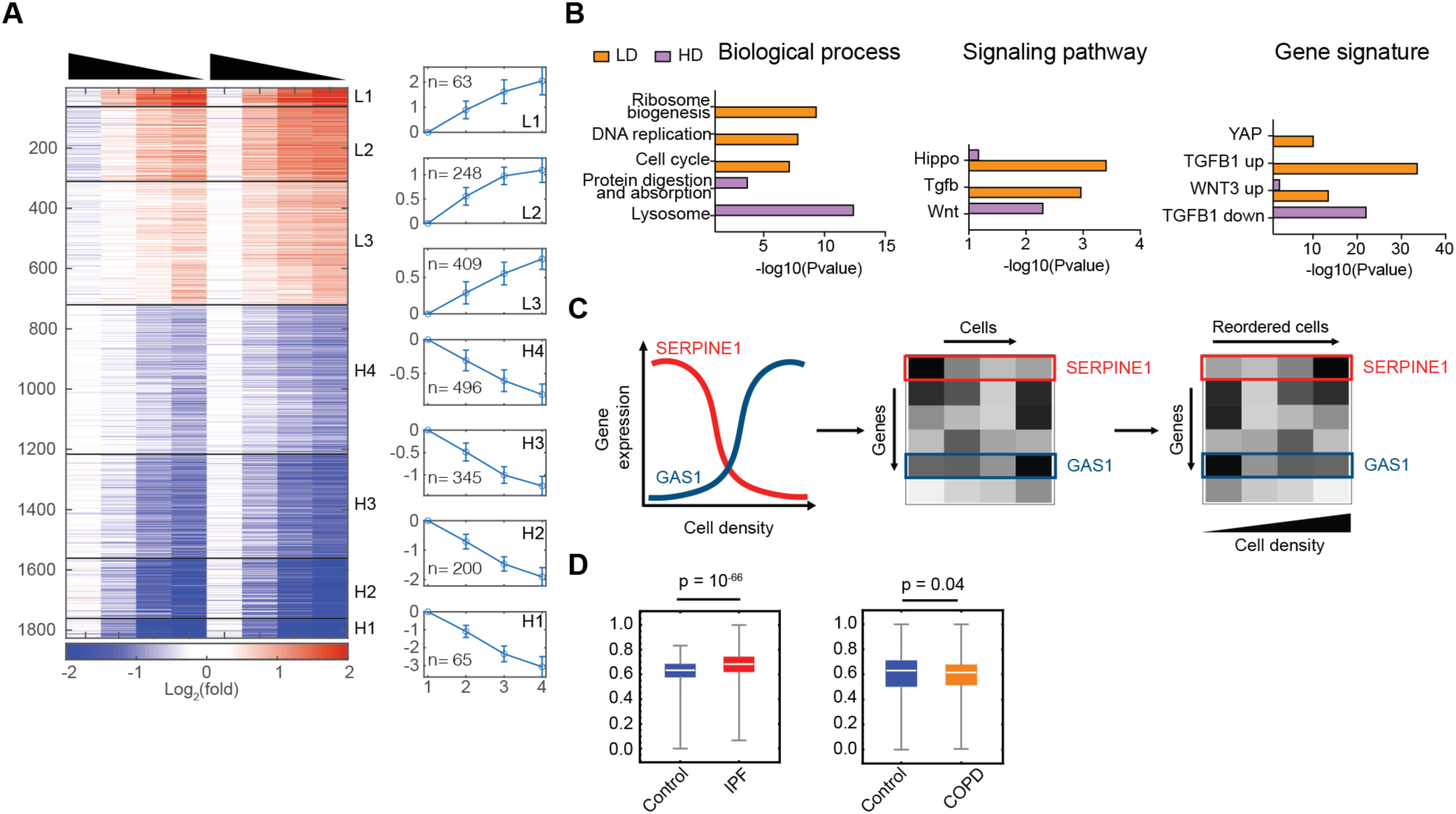
Fibroblasts exhibit density-dependent gene expression. A. Heatmap showing relative expression of density-dependent genes in MEFs from two sets of biological replicates. Gene expression (TPM) is normalized to the average of two replicates at 10K density and shown after log2 transformation. Genes are clustered into 7 groups using unsupervised K-mean clustering, organized from the most induced at low density to the most induced at high cell density. The average log2 fold change of each cluster and the number of genes is shown on the right. Density of cells is indicated by the triangle above the heatmap, high to low. B. Functional enrichment analyses of biological processes, KEGG signaling pathways, and molecular signatures for genes induced in low density (LD) or high density (HD). C. Using landmark genes that are density-dependent, cells in single-cell gene expression data can be reordered along a pseudo density axis to find new density-dependent gene programs. D. Violin plots showing the distribution of calculated cell density scores for ScRNAseq of human fibroblasts extracted from IPF+control and COPD+control groups.

To understand the functions of the density-dependent expression programs, we performed enrichment analysis for these clusters independently and in combination. Genes induced at low cell density are enriched for ribosome biogenesis, cell cycle, DNA replication, and purine and pyrimidine metabolism, consistent with higher anabolic activity of proliferating cells at low cell density. In contrast, genes upregulated at high cell density are enriched for genes associated with lysosomal function, extracellular matrix, and protein digestion and absorption (Figure 2B, Figure S1D). Signaling pathways, including MAP kinase signaling, Hippo-YAP signaling, TGF-*β* signaling, Ras, VEGF, TNF and IL-17 signaling pathways are significantly enriched at low cell density, and Wnt signaling pathway is enriched at high cell density (Figure 2B, Figure S1E). To further test if these signaling pathways may regulate the cell density-dependent expression programs, we examined the enrichment of gene expression signatures curated in the Molecular Signature database among density-dependent clusters (Liberzon et al., 2015; Subramanian et al., 2005). Hippo-YAP, TGF-*β*, and Wnt activation demonstrate the strongest enrichment for genes induced at low cell density, and genes repressed by TGF-*β* show the strongest enrichment at high cell density (Figure 2B, Figure S1F). Overall, functional enrichment analysis revealed that Hippo-YAP and TGF-*β* signaling pathways are most highly correlated with density-dependent gene expression.

Space limitation *in vitro* in tissue culture dishes is distinct from space limitation within tissues due to the complexity of tissue microenvironment (Duval et al., 2017). We sought to test if the density-dependent expression programs we identified *in vitro* suggest similar mechanisms of cell density sensing in physiological or pathological conditions. Fibrosis is characterized by excessive fibroblast proliferation (Wynn, 2008). Recently, single cell data of lung fibrosis samples from humans have become available (Adams et al., 2020). Inspired by the previous work that reconstructed the spatial environment of single cells based on gradient expression (Halpern et al., 2017; Moor et al., 2018), we developed a computational algorithm to estimate ‘cell density’ for individual cells based on their relative expression of density-dependent genes (Figure 2C). This allowed us to compare the predicted cell density of fibroblasts between healthy individuals and patients with idiopathic pulmonary fibrosis (IPF) or chronic obstructive pulmonary disease (COPD) (Neumark et al., 2020). Intriguingly, fibroblasts isolated from IPF patients display an increased tendency for high cell density (Figure 2D, S2A), consistent with the known invasive expansion of myofibroblasts (Scotton and Chambers, 2007). On the other hand, fibroblasts from COPD patients seem to be largely similar to healthy individuals (Figure 2D, S2B). These results suggest that the density-dependent expression programs identified *in vitro* may represent changes in fibroblasts that occur during fibrotic disease.

### Space availability modulates the Hippo-YAP and TGF-β signaling pathways

The Hippo signaling pathway can be activated by diverse cellular and environmental signals and converges on a pair of homologous transcription factors YAP1 and TAZ (Varelas, 2014). When appropriate environmental signals are present, such as cell-cell contact, YAP1 is excluded from the nucleus and either sequestered or degraded. When Hippo signaling is off, YAP1 translocates into the nucleus, where it forms a complex with the TEAD family DNA-binding transcription factors to regulate target gene expression. TGF-*β* signaling is activated by TGF-*β* family members, often bound to ECM in a latent form, that become activated through protease, integrin, or other processing events (Li and Flavell, 2008). Activation of TGF-*β* receptors leads to phosphorylation of receptor-regulated SMADs (R-SMADs), particularly SMAD2 and SMAD3. Phosphorylated R-SMADs translocate into the nucleus, forming a complex with SMAD4 to regulate target gene expression (Macias et al., 2015)

We found that both TEAD and SMAD binding sequence motifs are significantly enriched in the promoters of genes upregulated at low cell density (Figure 3A), while only SMAD binding motifs are enriched for genes induced at high cell density (Figure 3A). SMAD proteins can act as both transcriptional activators and repressors (Massagué et al., 2005). Distinct SMAD motifs enriched among the genes induced at either high or low densities suggest that cooperation with other transcriptional co-activators or co-repressors may determine density-dependent activation or repression in addition to TGF-*β* signaling.

**Figure 3.**
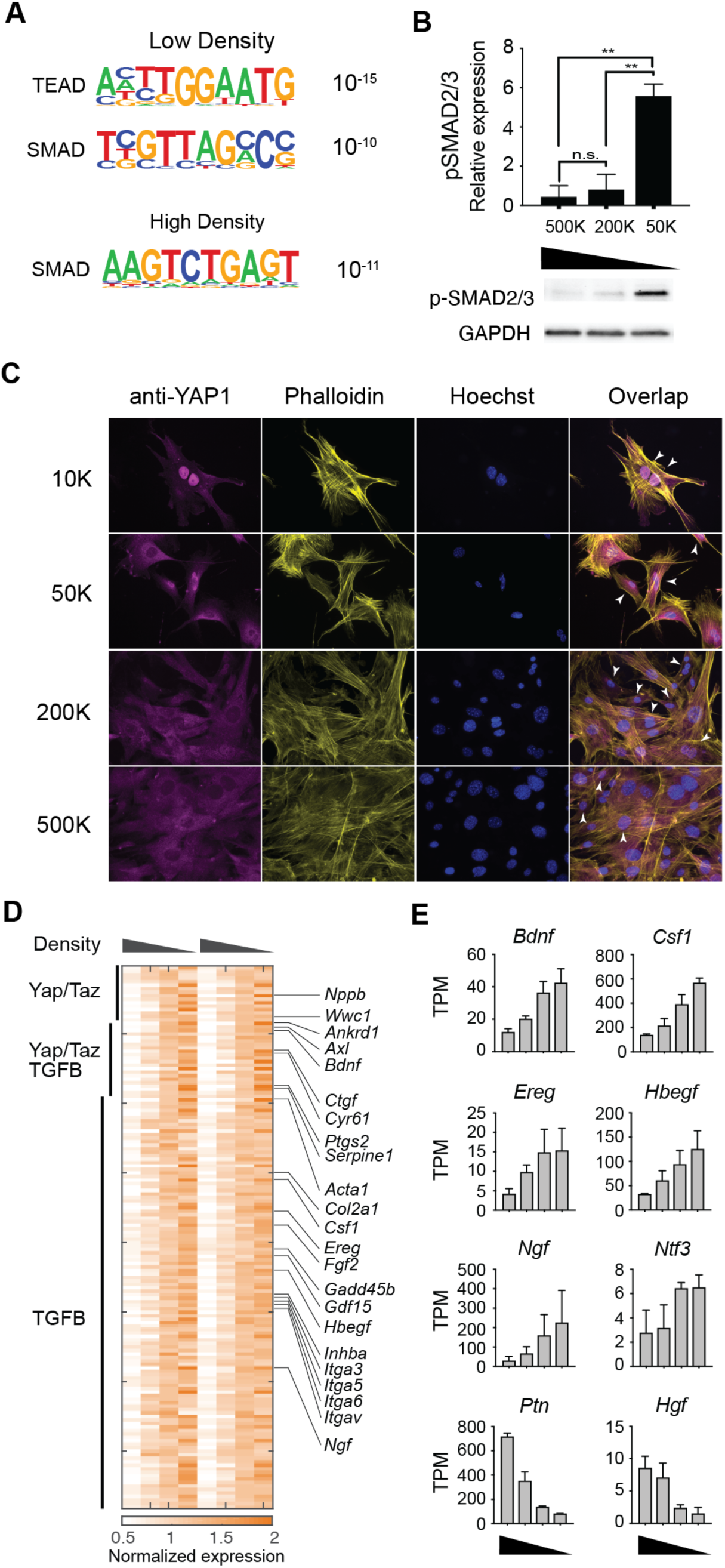
YAP1 and SMADs are activated in a density-dependent manner in fibroblasts. A. Significantly enriched transcription factor binding motifs in genes induced at low or high density. B. Phosphorylation of SMAD2/3 at different cell densities. Top, quantification of phospho-Smad2/3 Western blot staining, normalized to GAPDH staining of the same sample, from 3 independent experiments. Bottom, representative image of phospho-Smad2/3 and GAPDH Western blots. C. Representative immunofluorescent images showing YAP1 localization in MEFs cultured at the specified cell densities overnight. Arrowheads denote examples of nuclear localization of YAP1. D. Heatmap showing annotated targets of Hippo-YAP signaling and TGF-*β* signaling pathways at different cell densities. E. Expression of representative growth factors significantly regulated by density (high to low density).

Next, we wanted to confirm whether the activities of the Hippo-YAP and TGF-*β* signaling pathways are regulated by cell density. Significantly increased phosphorylation of SMAD2/3 was observed at low cell density, suggesting that activation of SMAD proteins contribute to the observed density-dependent transcription programs (Figure 3B & S3A). To examine the role of Hippo-YAP signaling in density-dependent gene expression programs, nuclear localization of YAP1 was used to infer activity of the Hippo-YAP pathway. As expected, YAP1 is localized to the nucleus at low cell density. As cell density increases, YAP1 is gradually excluded from the nucleus, as indicated by the decrease in nuclear staining (Figure 3C). Collectively, these data demonstrate that the activity of both TGF-*β* signaling and Hippo-YAP signaling are regulated in fibroblasts in a cell density-dependent manner, in response to space limitation. Indeed, several genes that are known to be directly controlled by YAP1/TAZ and SMAD proteins display density-dependent gene expression in fibroblasts. These genes include YAP1/TAZ targets, such as *Nppb*, *Akred1*, *Bdnf*, *Ctgf*, *Cyr61*, as well as TGF-*β* target genes, such as *Serpine1*, *Acta1*, *Col2a1*, *Hbegf*, *Ngf* (Figure 3D).

### YAP1 and TGF-β signaling control expression of different growth factors in response to space limitation

Within the group of genes regulated by space limitation, we observed several growth factors specific for different cell lineages (Figure S3B). For example, neurotrophic growth factors (*Bdnf, Ngf, Nif3, Ptn*), epidermal growth factors (*Ereg, Hbegf*), hepatocyte growth factor *Hgf*, and myeloid growth factor *Csf1*, all show density-dependent expression patterns (Figure 3E). In particular, the expression of *Csf1* was inversely related to fibroblast density, suggesting that density sensing by fibroblasts is coupled with *Csf1* production for macrophages (Zhou et al., 2018). To examine which pathway controls the expression of *Csf1* in fibroblasts, we tested whether activation of YAP1, TGF-*β*, or Wnt signaling is sufficient to regulate *Csf1* expression. Recombinant TGF-*β* or Wnt3a do not induce the expression of *Csf1* at the mRNA level (Figure 4A). However, *Hbegf* and *Ctgf* are induced by TGF-*β*, and to a lesser degree by Wnt3a (Figure S3C). Next, we tested if activation of YAP1 can control *Csf1* expression. To activate YAP1 in primary fibroblasts, we used a previously established genetic model with constitutively active YAP1 (Su et al., 2015). A mutation from Serine to Alanine prevents phosphorylation at residue 112 of YAP1 and results in constitutive nuclear localization. Overexpression of a similar mutant in the murine liver causes an increase in liver size, consistent with the known role of YAP1 in organ size control (Camargo et al., 2007). In our model, *Yap1^S112A^-IRES-GFP* is expressed at the *Rosa26* locus, downstream of a floxed transcriptional STOP cassette. After introducing MSCV containing Cre recombinase, primary fibroblasts carrying the *Yap1^S112A^* allele express constitutively active YAP1 (referred to as YAP1^CA^). We found that the expression of *Csf1* is significantly elevated in these cells, in comparison to primary fibroblasts transduced with MSCV carrying GFP alone (Figure 4A).

**Figure 4.**
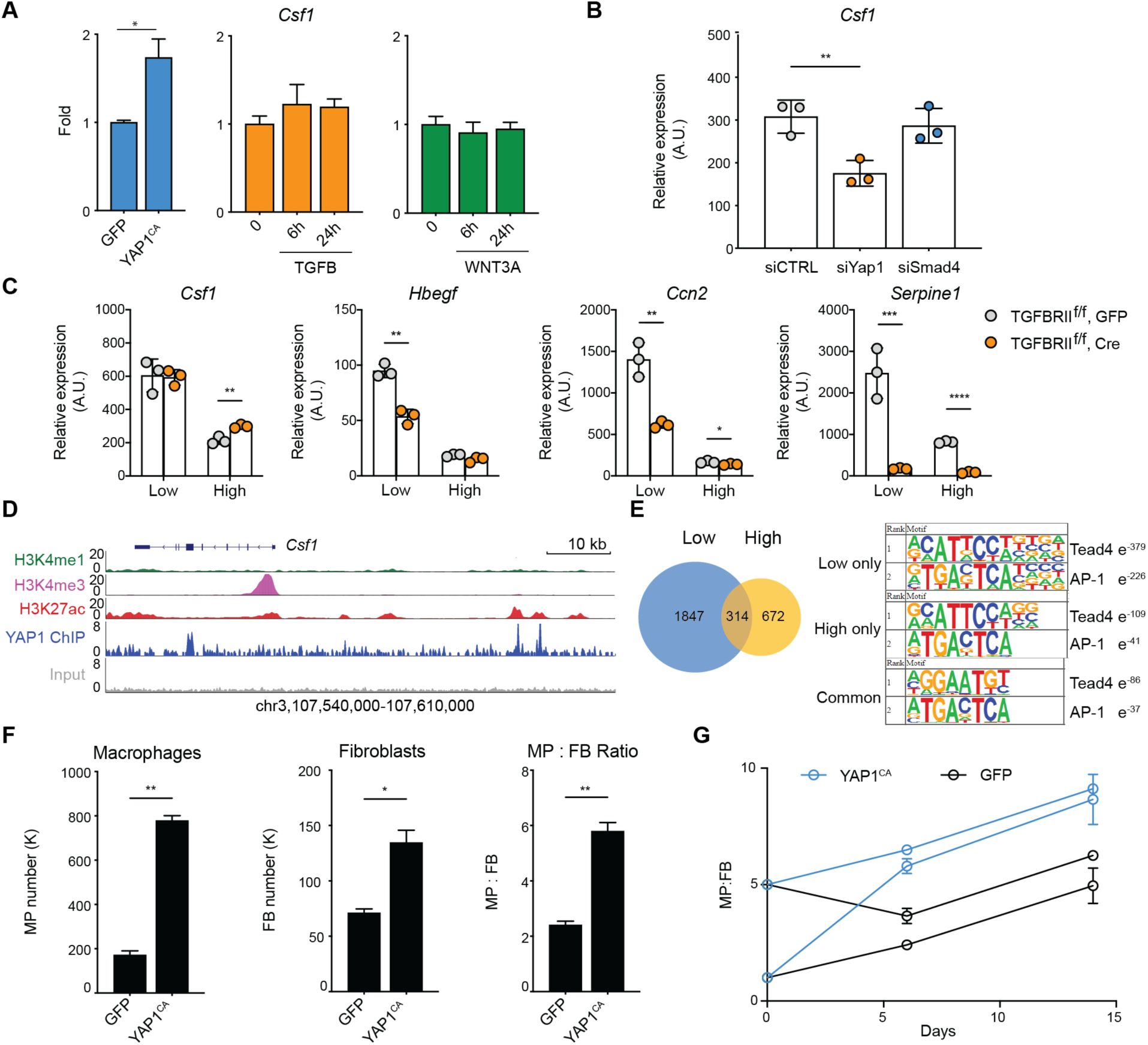
YAP1-dependent regulation of *Csf1* in fibroblasts controls macrophage numbers. A. *Csf1* expression in MEFs carrying Yap^CA^ (left), or treated with recombinant TGF-*β* (center) or WNT3A (right). MEFs were isolated from YapKI^fl/fl^ mice and then transduced with lentivirus carrying GFP (GFP) or Cre-GFP (constitutively-active, YAP^CA^). B. *Csf1* expression in MEFs 3 days after transfection with siRNAs targeting *Yap1*, *Smad4,* or scrambled control siRNA (siCTRL). C. Expression of selected genes in wild type and *Tgfbr2* KO MEFs cultured at low or high density. Wild type and *Tgfbr2* KO MEFs are generated by viral transduction of *Tgfbr2*^fl/fl^ MEFs with GFP or Cre-GFP vectors respectively. D. Genomic tracks displaying ChIP-seq occupancy of histone modifications (H3K27ac, H3K4me1, and H3K4me3) and endogenous YAP1 binding at the *Csf1* gene locus in MEFs. E. Venn diagram showing the number of YAP1 binding peaks at low and high cell densities determined by ChIPseq. The top ranked enrichment of transcription factor motifs is shown for peaks at high, low, or both cell densities. F-G. Wild type (GFP) and Yap^CA^ MEFs were plated together with WT BMDMs. Their numbers and ratios were determined by flow cytometry 6 days after co-culture (F). BMDMs & MEFs were plated at different starting ratios and quantified at day 6 and 14 (G).

These data suggest that *Csf1* is induced by the activation of YAP1, but not TGF-*β* or Wnt signaling. However, overexpression of a constitutively active transcription factor may not faithfully reflect the role of endogenous YAP1 in response to space limitation. Therefore, we used siRNA knock-down to validate the functions of endogenous YAP1. Indeed, knocking-down *Yap1* reduced *Csf1* expression by about 2-fold, similar to the difference between low and high cell densities, while knock-down of *Smad4*, the central transcription factor in TGF-*β* signaling, had minimal effects (Figure 4B). Conversely, siRNA knockdown of *NF2*, an upstream suppressor of YAP1/TAZ, resulted in increased *Csf1* expression (Figure S3D). These data demonstrate that activation of endogenous YAP1 is necessary and sufficient to control *Csf1*. Both YAP1 and TAZ are transcriptional activators downstream of Hippo-YAP signaling. They interact with TEAD transcription factors, bind to similar DNA sequence motifs, and are both expressed in fibroblasts (Figure S3E). Furthermore, siRNA against *Wwrt1* (the gene encoding TAZ protein) had a partial effect on *Csf1* expression but targeting *Wwrt1* and *Yap1* simultaneously had no additive effect compared to targeting *Yap1* alone (Figure S3D). This suggests that YAP1 is the primary transcriptional activator controlling *Csf1* expression. However, it is worth noting that YAP1 and TAZ have differential effects on controlling the expression of other genes (data not shown). These results are not due to differences in siRNA targeting efficiency, as 90% knock-down was achieved for all targets at the level of mRNA expression (Figure S3F). Finally, using *YAP1^CA^* fibroblasts, we compared gene expression of a subset of density-dependent growth factors to WT fibroblasts and identified additional growth factors regulated by YAP1 activity (Figure S3G). Expression of these growth factors also depends on endogenous levels of YAP1, as siRNA knockdown of *Yap1* reduces their expression (Figure S3H).

To further evaluate the TGF-*β* pathway in density-dependent growth factor expression, we used genetic and pharmacological targeting strategies. To genetically target TGF-*β* signaling, we isolated fibroblasts from *Tgfbr2*^fl/fl^ mice (Levéen et al., 2002), transduced them with either Cre-GFP or GFP viral vectors, and sorted them for GFP-positivity to indicate successful transduction.

In these cells deficient for TGFBR2, the main signaling receptor for TGF-*β* family ligands, expression of *Csf1* was unaffected (Figure 4C). In contrast, *Hbegf*, *Ctgf*, and *Serpine1*, genes regulated by cell density and TGF-*β* signaling, were significantly reduced or entirely abolished (Figure 4C). Similarly, pharmacological inhibition of TGF-*β* receptor achieved similar results (Figure S3I). Using both genetic and pharmacological approaches, we demonstrated that signaling through TGFBR2 is responsible for density-dependent regulation of a subset of growth factors including *Hbegf* and *Ctgf.* In contrast, the expression of *Csf1* was found to be controlled by YAP1, independent of TGF-*β* signaling.

### YAP1 regulates the expression of *Csf1* via a conserved distal enhancer

We next asked how the Hippo pathway regulates *Csf1* expression. *Csf1* is not known to be a YAP1 target gene and its promoter lacks a binding sequence for TEAD transcription factors. We speculated that YAP1 may regulate the expression of *Csf1* through distal regulatory elements. To test this hypothesis, we performed ChIP sequencing of endogenous YAP1 proteins in fibroblasts and discovered two distinct binding sites at 33 and 36 kb upstream of the *Csf1* transcription start site (Figure 4D). Analysis of histone modifications from the mouse ENCODE project (Davis et al., 2018; Dunham et al., 2012) indicated that both YAP1 binding peaks are within regions that are enriched for H3K27ac but lack H3K4me1 and H3K4me3 marks, suggesting that YAP1 physically occupies two active distal enhancers of *Csf1* (Figure 4D). On the other hand, we observed SMAD3 binding coinciding with H3K27Ac at the *Hbegf* locus in fibroblasts (Ruetz et al., 2017), supporting both published data and our observations that *Hbegf* is induced downstream of TGF-*β* (Figure S4A). At the global level, we observed about three times more YAP1 binding events at low density compared to YAP1 binding events at high density (Figure 4E). Both TEAD4 and AP-1 motifs were identified as highly enriched in YAP1 peaks, yet this enrichment was more robust at low density (Figure 4E). This is consistent with previous work showing that YAP/TAZ, TEADs, and AP-1 family members cooperate to regulate gene expression (Zanconato et al., 2015). Near the center of the +30 kb YAP1 peak at *Csf1* gene, we identified two sequences matching the consensus binding motifs of TEADs. Genomic sequence alignment of this region demonstrated high conservation of these sites between a variety of mammalian species, including opossum, dog, rat, rhesus macaque, and human (Figure S4B). These data suggest that the mechanism by which Hippo-YAP signaling regulates *Csf1* expression may be conserved in mammals. Based on H3K27ac data from the human ENCODE project (Davis et al., 2018; Dunham et al., 2012), we identified a similar enhancer ∼30 kb upstream of the human *Csf1* gene that contains the conserved TEAD binding sites. Interestingly, the activity of this 30 kb enhancer is cell type specific, in that the H3K27Ac mark is observed in fibroblasts, but not in endothelial or myeloid cells (Figure S4C). Conversely, an enhancer ∼12 kb upstream of the transcription start site, which has the AP1, but lacks the TEAD binding motifs, was enriched for H3K27ac in endothelial cells and myeloid cells, but not in fibroblasts (Figure S4C). These data suggest that *Csf1* expression is regulated by cell type-specific enhancers, and that density-dependent control of *Csf1* by YAP1 is a feature of fibroblasts.

### Fibroblasts produce CSF1 to support macrophage populations

Production of CSF1 from local tissues is essential for the maintenance, proliferation, and differentiation of tissue-resident macrophages (Buechler et al., 2021; Guilliams et al., 2020). Our previous work has demonstrated that fibroblasts produce CSF1 to support growth and survival of macrophages (Zhou et al., 2018). Similar communication has been observed between fibroblasts cells and tissue-resident macrophages *in vivo* (Bellomo et al., 2020; Bonnardel et al., 2019; Mondor et al., 2019). These communication circuits are essential to maintain homeostasis of macrophage numbers. By modeling the paracrine communication between macrophages and fibroblasts (Adler et al., 2018, 2020), we predict that changes in the expression of *Csf1* in fibroblasts affect the homeostatic number of macrophages as well as the relative ratio between macrophages and fibroblasts (Figure S4D). To test whether fibroblasts are a physiologically relevant source of CSF1 for macrophages *in vivo*, we used a genetic model of CSF1-deficiency in PDGFRα^+^ cells, including various fibroblast populations such as hepatic stellate cells. In the livers of PDGFRa^Cre^*Csf1*^fl/fl^ mice, we observed a decrease in macrophage frequency both by flow cytometry and histological analysis (Figure S4E, S4F). These data demonstrated that fibroblast populations provide an important source of CSF1 for macrophages *in vivo*.

In order to determine whether YAP1 regulation of *Csf1* expression in fibroblasts can directly control macrophage numbers, we returned to our previously established macrophage-fibroblast *in vitro* co-culture system in combination with YAP1^CA^ fibroblasts. As expected, constitutive activation of YAP1 resulted in an increase in fibroblast numbers, consistent with a role for Hippo-YAP signaling in autonomous control of cell proliferation (Figure 4F). In addition, macrophage numbers, as well as the overall macrophage to fibroblast ratio, significantly increased when YAP1 was active (Figure 4F). With elevated expression of *Csf1*, macrophages and fibroblasts still exhibited a stable ratio regardless of starting conditions (Figure 4G). This feature of stability was consistent with our previous observations and theoretical model prediction. It also demonstrated that YAP1-mediated regulation of *Csf1* is sufficient to control the number of macrophages in a defined compartment.

Like most growth factors, CSF1 acts in the vicinity of its source, thereby affecting both local numbers and spatial arrangement of macrophages (Buechler et al., 2021). We employed agent-based modeling to gain insights into whether density-dependent CSF1 production impacts the spatial distribution of macrophages. Agent-based modeling is a computational approach for simulating the behavior of multiple agents (cells in this case) that interact with each other according to specified rules (Glen et al., 2019; Wilensky and Rand, 2013). In the simulations, macrophages and fibroblasts are represented as red and blue cells respectively, and obey the following rules: (1) red cells produce a paracrine growth signal (*B1*, e.g. PDGF*β*) for blue cells, blue cells produce a paracrine growth signal (*R*, e.g. CSF1) for red cells and an autocrine growth signal (*B2*, e.g. PDGF*α*/HBEGF) (Zhou et al., 2018), (3) every cell computes the total amount of growth factor in its vicinity (tunable parameter), and undergoes one of three fate choices: die, stay, or divide, depending on the amount of available growth factor. The threshold values for cell death, survival, and proliferation are tunable parameters of the system. To model the density-dependent regulation of CSF1, we considered that the production of signal *R* from a blue cell is a decreasing function of the overall blue cell density in its vicinity. This model recapitulates the bi-stability of the two-cell circuit (Adler et al., 2018; Zhou et al., 2018), where the steady-state of the system depends on the initial conditions of these two populations (Fig. 5B-C). However, the impact of density-dependent regulation on the spatial distribution of red and blue cells is mild and sensitive to the choice of parameter values (Figure S5A-D). Tissues are often composed of much more than two cell types. To test if the density-dependent regulation may have an oversized role in conditions of complex cell composition, we introduced a third ‘inert’ cell type (or a non-cell structure) that takes up space but does not interact with either blue or red cells via growth factors (Fig 5A, lime cell). We considered several arbitrary structures of lime cells and presented the simulation where lime cells are organized as rings of different radius. Without density-dependence, the steady-state spatial pattern of red and blue cells is uniformly distributed throughout (Fig 5D). Interestingly, with density-dependence, red cells localize away from the lime cells, as if the red cells are excluded by the lime cells. This effect is even more striking in the center of small lime cell rings, where the loss of red cells impacts the survival of accompanied blue cells, eventually creating an empty zone (Fig 5E, S5E-H). This may represent conditions where the emergence of non-cellular structure or growth of ‘inert’ cells exclude macrophages and fibroblasts (such as in late-stage fibrosis). Overall, agent-based modeling of macrophages and fibroblasts revealed an intriguing role of density-dependent production of paracrine growth factors in regulating macrophages in space, which may impact the spatial organization of macrophages in complex tissues.

**Figure 5.**
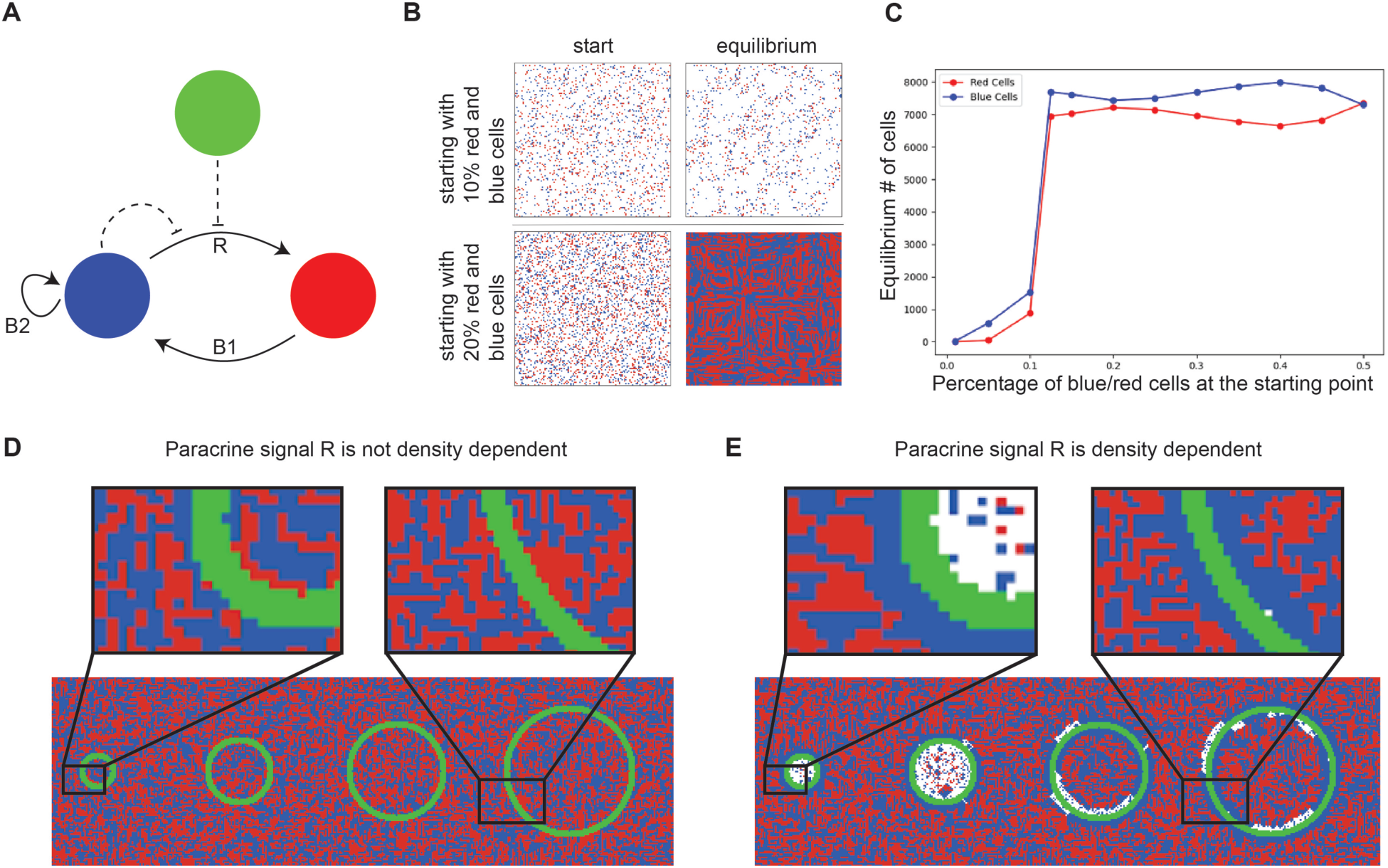
Agent-based model of the macrophage-fibroblast cell circuit reveals impact on spatial organization. A. Schematics of agent-based modeling describing the interactions between MPs and FBs. Blue cells, FBs; red cells, MPs; lime cells, “inert” cells. B. Representative simulation of the two-cell circuit in 2D. Two snapshots were taken at the start and at equilibrium for each simulation, at initial conditions with low cell density (upper panels) and high cell density (lower panels). C. Graph showing the steady-state cell number as function of the initial density of blue and red cells. D, E. Representative images of simulation with a third cell type, without (D) and with (E) density-dependent regulation of paracrine growth factor *R*. Lime, inert cells or non-cell structure.

### Space limitation regulates gene expression through actin-dependent mechanisms

The Hippo-YAP and TGF-*β* pathways are well known to be involved in sensing cellular environment (Duronio and Xiong, 2013; Varelas et al., 2010). YAP activity can be regulated by mechanical properties and composition of ECM. TGF-*β* signaling can be induced by soluble and contact-dependent signals. How these signaling pathways are induced by space availability remains elusive. We first tested whether the density-dependent expression programs in fibroblasts were due to soluble signals or responses to ECM components. We transferred supernatants of low-density cultures to high-density cells, and vice versa, to test whether fibroblasts sense density signals through soluble factors (Figure S6A). We also transferred cells onto decellularized ECM, from low- or high-density cells, to test whether fibroblasts sense density through properties of ECM (Figure S6B). However, neither supernatant transfer nor ECM “transfer” mimicked the effect of either low- or high-density conditions. Although these experiments do not exclude the role of ECM, they suggest that fibroblasts may sense cell density or space availability through additional, cell-intrinsic mechanisms.

We observed that the average surface area of a cell is smaller at high density than that of a cell at low density (Figure S6C, D). The length and width of nuclei are also smaller at high cell density than at low cell density (Figure S6E, F), while the height or ‘thickness’ of nuclei shows the opposite trend (Figure S6G), and similar nuclear volume is maintained across conditions (Figure S6H). We reasoned that lack of contact with neighboring cells allows expansion of the cell body and thus flattens the nucleus (Figure 6A). To examine the change in nuclear shape, we quantified the sphericity of nuclei using the ratio between the length and the width (the longest and second longest axes of nuclei respectively). A spherical nucleus has a ratio close to one while a flattened nucleus has a much smaller ratio. Indeed, nuclei in cells at low cell density are significantly less spherical than in cells at high density (Figure 6B), indicating that nuclei at low and high cell densities experience different mechanical pressure.

**Figure 6.**
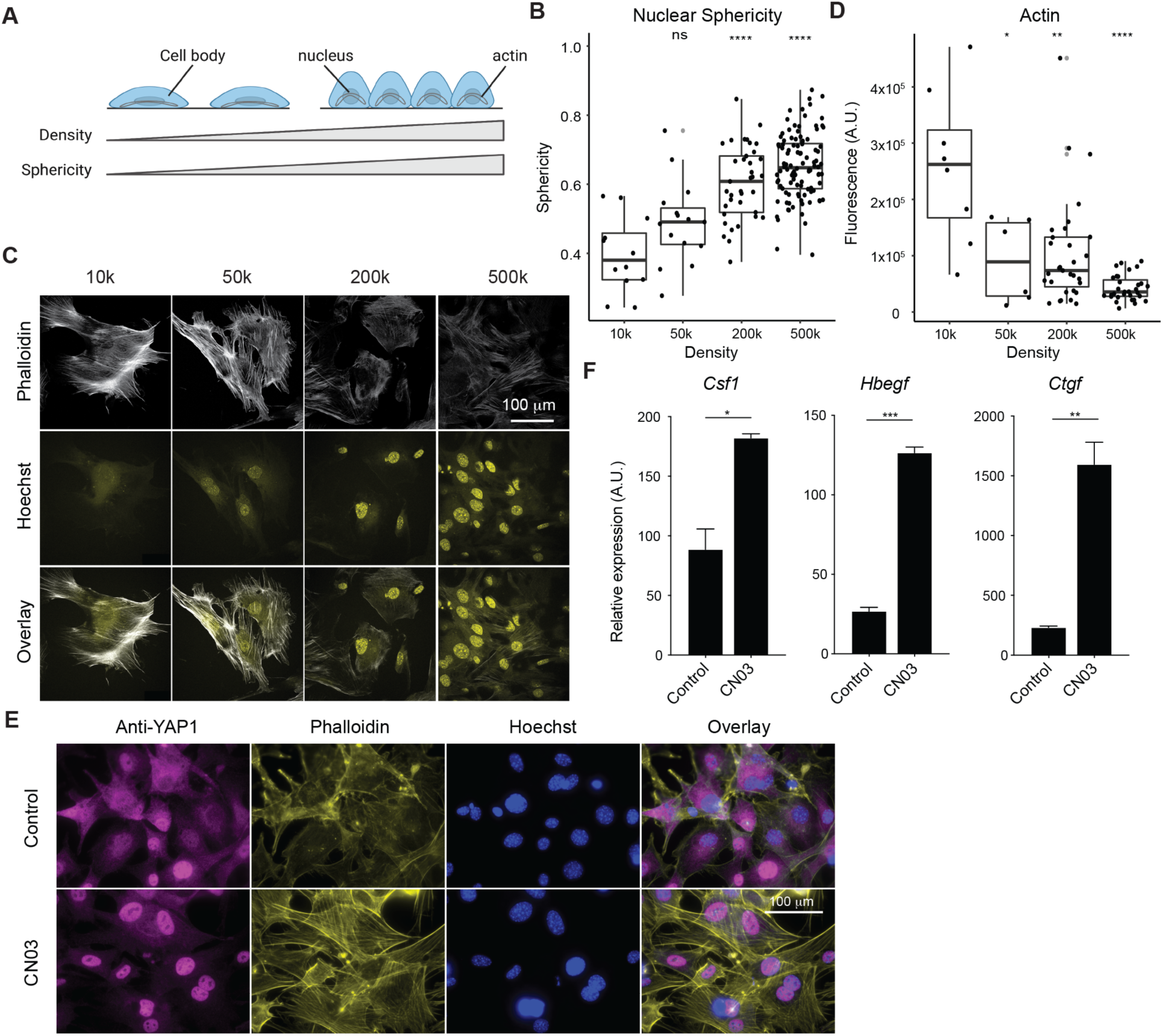
Actin-dependent mechanisms regulate density gene expression. A. Diagram depicting the relationship between increased nuclear sphericity and increased cell density. B. Quantification of nuclear sphericity of MEFs, calculated as the ratio between the length and width of a nucleus. Each point represents an individual nucleus. Six images per density were quantified. Wilcox t test. ns p>0.05, * p<0.05, ** p<0.01, *** p<0.001, **** p<0.0001. C. Confocal images of immunofluorescent staining of actin filaments (Phalloidin) in MEFs at different cell densities. D. Quantification of fluorescence intensity of actin staining at different cell densities. Fluorescence intensity of actin inside the cell is corrected by the background. Each data point represents one cell. E. Representative immunofluorescence images of YAP1 localization in MEFs after 4 hour treatment with CN03. F. Growth factor expression in MEFs after 4 hour treatment with Rho activator (CN03 peptide).

Previously, it was reported that stiff matrices promote nucleation of actin fibers that apply pressure to the nucleus, leading to YAP1 nuclear translocation (Elosegui-Artola et al., 2017; Shiu et al., 2018). In our experiments, discrete actin fibers are commonly observed at low density (Figure 6C) and display higher fluorescent intensity (Figure 6D). We thus tested if formation of actin fibers directly regulates the expression of density-dependent growth factors. CN03, a peptide derived from bacterial deamidase toxins, activates RhoA by locking it into constitutively active form (Schmidt et al., 1997). RhoA then activates Rho activated protein kinase (ROCK), which promotes actin polymerization through LIM kinase and coffilin, and contractility of actin fibers through activation of myosin-light chain kinase and inhibition of myosin-light chain phosphatase (van Aelst and D’Souza-Schorey, 1997; Arber et al., 1998; Yang et al., 1998). Following two hours of treatment with CN03, we observed that actin filaments formed across the length of cells, and that YAP1 was found almost exclusively in the nucleus even at high cell density (Figure 6E). Interestingly, activation of RhoA strongly induced the expression *Csf1* and other YAP1-dependent growth factors including *Bdnf* and *Ereg* (Figure 4F, Figure S7A), and siRNA knock-down of *Yap1* abolished the RhoA-dependent activation of these genes (Figure S7A). To further test if actin assembly is responsible for elevated *Csf1* expression when cells grow at low density, we treated low density cells with either Y27632 (Uehata et al., 1997), a ROCK inhibitor, or Latrunculin A (Spector et al., 1989), an inhibitor of actin assembly. We found that blocking the formation of actin filaments was sufficient to reduce nuclear localization of YAP1 and expression of *Csf1* (Figure S7B, C). In addition to YAP1-regulated growth factors, two growth factors that are strongly dependent on cell density and TGF-*β* signaling, *Hbegf* and *Ctgf,* are also activated by RhoA, but independent of YAP1 (Figure 6E, Figure S7A). Indeed, treatment of CN03 triggers significant enrichment of SMAD2/3 in the nucleus, and this localization can be reversed by inhibiting actin assembly with Y27632 (Figure S7D). Both Y27632 and Latrunculin A can reduce the expression of *Ctgf* and *Hbegf* in cells cultured at low density (Figure S7B). Moreover, at the transcriptomic level, density-dependent expression programs are highly enriched for genes that are regulated by Rho signaling (MsigDB, “Berenjeno transformed by RhoA”, p < 10^-114^). Altogether, these data demonstrated that the activity of both YAP1 and SMAD can be regulated by actin assembly. Given the distinct change of nuclear shape at different cell densities, these data suggest that sensing space availability may be dependent on RhoA signaling, and that density-dependent expression programs may be controlled through mechanical forces acting on the nucleus.

### Environmental conditions regulate growth factor expression

In ecology, extrinsic environmental factors determine the carrying capacity, or maximum size, of a given population. In a tissue compartment, space is one such limiting factor. We demonstrated that the cell type sensitive to space limitation regulates the expression of growth factors for a cell type that is insensitive to space constraints. A variety of other environmental variables, including nutrient or oxygen levels, could limit the number of cells in a compartment. We thus screened expression of growth factors in fibroblasts under diverse conditions, including amino acid deprivation, glucose deprivation, and hypoxia. Several growth factors are strongly induced or repressed by each condition (Figure 7A). To test if there is a pattern in the control of growth factors, we included additional environmental conditions that may have effects on carrying capacity such as oxidative, osmotic, and endoplasmic reticulum (ER) stress. We analyzed how changes in expression vary among different conditions in comparison to the overall change at the transcriptomic level. Surprisingly, transcriptional changes of growth factor genes were highly correlated among oxidative stress, ER stress, and glucose and glutamine deprivation (Figure 7B). In contrast, the overall cellular responses to these stress conditions and nutrient limitations show poor correlation (Figure 7B). These analyses suggested that a subset of growth factors is regulated by factors of cellular environment. For example, *Ctgf*, *Vegfa*, *Hbegf*, *Lif*, *Gdf15*, *Il11*, and *Areg* are among the growth factors that are commonly induced, while *Bmp4*, *Vegfd*, *Hgf*, *Pdgfd* are among the growth factors that are commonly repressed under these conditions (Figure 7C). Altogether, these data reveal a central role for environmental sensing by fibroblasts in the control of growth factor production.

**Figure 7.**
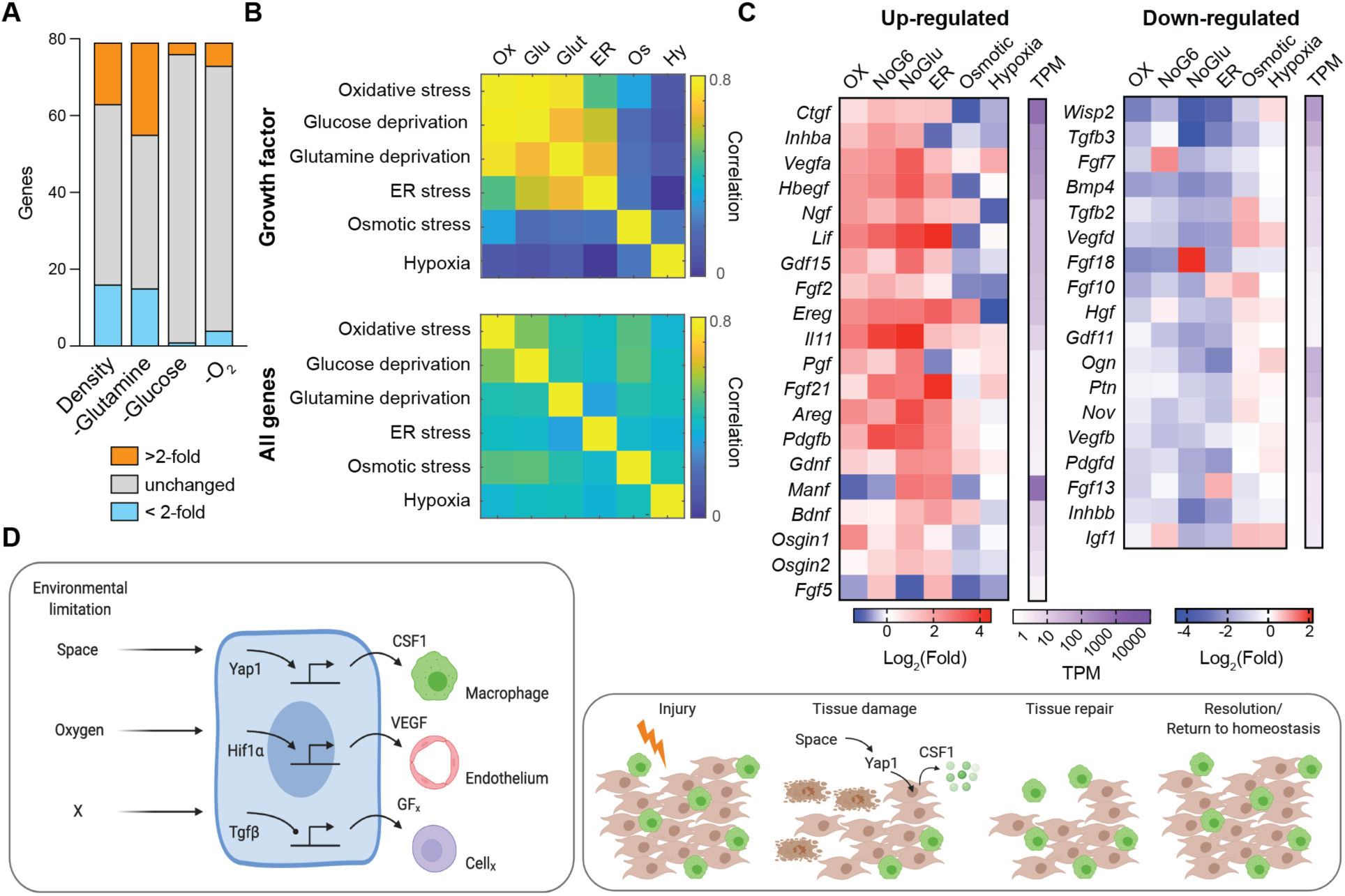
External environment regulates growth factor expression in fibroblasts. A. The number of growth factors differentially regulated in MEFs under different densities and nutrient-depleted conditions. B. Heatmap showing the correlation of growth factor genes or the complete transcriptome among different conditions. C. Heatmap showing expression fold changes of selected growth factors, grouped as commonly induced or repressed in MEFs at different conditions. D. Sensing of environmental conditions including space and oxygen regulates growth factor production through conserved signaling pathways such as Hippo-YAP (left). These mechanisms may explain how cell loss due to tissue damage is repaired and tissue homeostasis restored (right).

## Discussion

The population sizes of different cell types within a tissue compartment must be tightly regulated to ensure proper tissue functioning, prevent overgrowth, and allow for regeneration and repair (Figure 7D). Two modes of cell number control have been described previously: one is based on space availability, which can be mediated through either cell-cell or cell-ECM contact; the other is based on growth factor availability (Eagle and Levine, 1967; Humphrey et al., 2014; Raff, 1992). However, whether these control strategies function independently from each other is not known. Here we demonstrated that fibroblasts and macrophages, two universal cell types within tissues, each use a different strategy to control their numbers. Fibroblast proliferation is more sensitive to the availability of space, while macrophage expansion is highly sensitive to the availability of a growth factor. Moreover, we found that production of the macrophage-specific growth factor CSF1 by fibroblasts was directly regulated by Hippo-YAP signaling in response to space limitation. This coupling between space availability with growth factor production provides a simple link between the two different modes of cell number control (Figure 7D).

The Hippo-YAP pathway is known to regulate cell proliferation, apoptosis, stemness, and differentiation in response to a variety of internal and external signals (Camargo et al., 2007; Gumbiner and Kim, 2014; Su et al., 2015; Varelas, 2014; Yu et al., 2015). Therefore, its role in controlling cell density-dependent responses was not unexpected. However, its involvement in the regulation of growth factor expression was intriguing because this pathway has been most commonly described for cell autonomous (or cell-type autonomous) responses. However, in a recent study, YAP was found to not only inhibit pluripotency of stem cells autonomously, but to also induce pluripotency of neighboring cells through production of matrix proteins (Hartman et al., 2020). This work along with our findings suggest that Hippo-YAP1 signaling also has important functions in regulating cell non-autonomous responses to environmental cues through cellular communication. Moreover, we found that changes in cell and nuclear shape that occur in sparse versus dense environments may reflect mechanical pressure of the actin cytoskeleton on the nucleus, which regulates translocation of YAP1 into the nucleus. Interestingly, we found that manipulation of RhoA signaling, and assembly of actin can regulate density-dependent gene expression that is dependent on both YAP1 and SMAD. Recently, it has been shown that pressure on the nucleus regulates cell migration behavior (Lomakin et al., 2020a; Venturini et al., 2020a). Our work suggests that the nucleus can be a mechanical sensor for cell density and control both cell-autonomous and non-autonomous proliferation. This is consistent with recent findings demonstrating the role of the cell nucleus in mechanosensing (Kirby and Lammerding, 2018; Lomakin et al., 2020b; Venturini et al., 2020b). Future studies will need to define the intermediary steps between changes in cell shape, actin mechanics, and activation of signaling pathways such as YAP and TGF-*β*.

Signaling pathways that sense tissue microenvironment are often not specific to a particular cell type. For example, epithelial cells, fibroblasts, stem cells, and hepatocytes all employ Hippo-YAP1 signaling pathways to control their proliferation in response to cell-cell and cell-ECM interactions (Gumbiner and Kim, 2014; Su et al., 2015; Varelas, 2014; Yu et al., 2015). We found that YAP1 regulates the expression of *Csf1* via a conserved distal enhancer that is uniquely active in fibroblasts. This Hippo-YAP1-regulated enhancer thus couples fibroblast density with macrophage numbers. This observation suggests that certain cells within tissues may have specialized functions in regulating tissue composition in response to different environmental factors (Figure 7D). For instance, macrophages are well known to sense hypoxia through HIF1α and in turn secrete VEGFA to promote endothelial growth and vascularization (Shweiki et al., 1992). Skin macrophages sense hyper-osmolarity and produce VEGFC to enhance proliferation of lymphatic endothelial cells (MacHnik et al., 2009). This link between environmental sensing by one cell type and growth factor production for another cell type may be a general feature employed to regulate tissue composition. Cell type-specific regulatory elements of growth factor gene expression may provide molecular fingerprints to uncover the underlying links between tissue microenvironment and cell type composition.

Not all cell types respond to growth factors or environmental signals by increasing rates of proliferation. Indeed, cells can be categorized as labile, stable, or permanent, based on their proliferative capacity. Permanent cells, such as neurons and cardiomyocytes, have limited proliferative capacity and their numbers in adults are primarily determined during development. Labile cells, such as intestinal epithelial cells, skin keratinocytes, and most hematopoietic cell types, are continuously replenished from stem cells and have high proliferative capacity until they reach a terminally differentiated state, at which point they become postmitotic. The appropriate ratios and numbers of cells going through different developmental stages can be in principle determined based on negative feedback between differentiated cells and their progenitors (Liang et al., 2017; Lo et al., 2009). Finally, stable cells, such as macrophages, fibroblasts, and hepatocytes, are normally quiescent but can proliferate even in a fully differentiated state in response to specific growth factors (Hoyer et al., 2019; Michalopoulos, 2007; Wynn, 2008). Through studying macrophages and fibroblasts, we found that population size control of two different stable cell types is coordinated: the signaling pathway that limits proliferation of one cell type regulates expansion of the other cell type. Tissue-resident immune cells, including mast cells, macrophages, and innate lymphoid cells, are distinct ‘stable’ cell types and their numbers are likely regulated in a similar fashion, by signals from stromal cells such as fibroblasts.

In an ecosystem, carrying capacity is defined as the maximum population size that can be supported in a given environment (Gotelli, 2008). In tissues *in vivo* or cell cultures *in vitro*, many variables can regulate cellular proliferation and survival, but carrying capacity is determined by the variables that are the most limiting for growth. At standard culture conditions, we found that fibroblasts and macrophages use different strategies of cell number control. The mechanism used by fibroblasts to detect changes in space availability directly controls growth factor production for macrophages. Fibroblasts are well known to be regulated by growth factors such as PDGFs and EGFs. However, the compartment size for fibroblasts at steady state is primarily determined by space availability. Additionally, we found that changes in other environmental factors, such as oxygen or nutrient levels, regulated growth factor production in fibroblasts. In this way, unique environmental stimuli such as mechanical cues, oxygen, or nutrients determine the levels of growth factors produced by fibroblasts for macrophages and other cell types (Figure 7D). Whether this is a general mechanism of growth factor regulation in other cell types that respond to limitations in space remains to be determined.

Lastly, our findings may also have implications in the progression of ‘tissue-level’ diseases such as cancer and fibrosis. One conserved feature of tumors, regardless of anatomical location or tissue of origin, is dysregulation of cell composition and escape from normal growth control mechanisms. Increased YAP/TAZ activity in tumor cells is one of the mechanisms linked to cancer initiation and progression. In fact, YAP/TAZ activation is observed in a broad spectrum of human cancers, leading to tumor cell proliferation, survival, and invasion (Zanconato et al., 2016). Additionally, YAP/TAZ upregulation in cancer-associated fibroblasts is associated with high grade tumors and poor prognosis via effects on ECM stiffness (Calvo et al., 2013). Our data suggest there may be an additional tumor-promoting function of YAP1 activation, the upregulation of growth factors within the tumor microenvironment. Whether YAP1 activity in cancer-associated fibroblasts drives the production of CSF1 or other growth factors is an important open question. Similarly, YAP/TAZ hyperactivation is observed in both the epithelial and fibroblast tissue compartments of fibrotic lesions (Kim et al., 2019). This association with fibrosis has largely been attributed to YAP1-dependent transcription of ECM remodeling genes and feed-forward enhancement of ECM stiffness and contractile actin formation (Liu et al., 2015). Indeed, we found that actin polymerization promotes YAP1 nuclear translocation and downstream growth factor production. The extent to which YAP1 control of CSF1 also contributes to fibrotic disease is not yet known. Furthermore, a more detailed understanding of the upstream signals driving nuclear YAP1 will have important implications for the normalization of transformed and/or fibrotic tissues.

### Limitations and caveats

While this study focused on the proof-of-concept and molecular mechanisms of growth factor production, it remains to be determined how different factors of cell microenvironment regulate growth factor production in different cell types *in vivo*. An additional limitation is that it remains unclear whether cells sense mechanical properties of the environment (e.g., ECM stiffness), cell density, or both as primary indicators of space availability within tissues. Addressing these questions will require development of *in vivo* models that allow for evaluation of ECM properties as well as spatially resolved dynamic monitoring of growth factor expression.

## Acknowledgements

We thank current and former members of the Medzhitov lab for helpful discussions. This work was supported by Howard Hughes Medical Institute, the Blavatnik Family Foundation, the Scleroderma Research Foundation, and a grant from the NIH (1R01 AI144152-01). X.Z. was supported by the Jane Coffin Childs Memorial Fund postdoctoral fellowship. R.A.F. was supported by the Cancer Research Institute Donald Gogel postdoctoral fellowship. M.A. was supported by the Fulbright Scholar Fellowship, the Zuckerman STEM leadership program, the Israel National Postdoctoral Award Program for Advancing Women in Science, and the European Molecular Biology Organization (EMBO) Long Term Fellowship. M.L.M. was supported by the NIH MSTP Training Grant T32GM136651 and NHLBI F31 predoctoral fellowship HL139116-01A1.

## Materials and methods

### Mice

C57BL/6J (stock #000664), Pdgfra-cre (stock #013148), and Tgfbr2^fl/fl^ (stock #012603) mice were purchased from Jackson Laboratory. YapKI^fl/fl^ mice were generated previously (Su et al. 2015). Csf1^fl/fl^ mice were generously provided by Sherry Abboud Werner. All mice were maintained in a specific pathogen-free facility and animal experimentation was conducted in accordance with institutional guidelines.

### Cell culture and differentiation

Bone marrow-derived macrophages (BMDMs) were differentiated from whole bone marrow from female mice (8 to 12 weeks old) in the presence of L929-conditioned media. Femurs and tibias were removed from mice, flushed, and exposed to hypotonic lysis (ACK lysing buffer, ThermoFisher) to remove red blood cells. Cells were plated in macrophage growth media (MGM) overnight (RPMI + 2 mM L-glutamine, 1 mM sodium pyruvate, 10 mM HEPES, 200 U/mL penicillin/streptomycin, 10% FBS, and 30% L929-conditioned media). The next day (day 1), nonadherent cells were harvested, washed, and replated in 15 cm Petri dishes in 20 mL of MGM. On day 4, 15 mL of MGM was added to the plate. On Day 6-7, cells were lifted with 3 mM cold EDTA in PBS for 15 min and plated at appropriate concentrations for coculturing. All cell cultures were maintained in a 37°C incubator at 5% CO2.

Mouse embryonic fibroblasts (MEFs) were harvested from male and female E13.5-E14.5 embryos and sorted for purity. Staged embryos were removed from a pregnant female by removing the uterus and separating each embryo from its amniotic sac. The head and ‘‘red tissue,’’ including fetal liver, were removed and discarded. If genotyping was required, the head served as the source of DNA and embryos were kept separated. The remaining portion of each embryo was minced using razor blades in 0.05% trypsin + EDTA and placed in a 37°C incubator for 30 min. After digestion, the tissue was transferred into a conical tube, washed with complete DMEM (DMEM + 2 mM L-glutamine, 1 mM sodium pyruvate, 10 mM HEPES, 200 U/mL penicillin/streptomycin, 10% FBS; GIBCO) and resuspended in complete DMEM in 15 cm tissue culture plates overnight. The following day, cells and undigested tissue debris were lifted from the plates using 0.05% trypsin + EDTA, spun down, resuspended, and filtered over a 70 μm filter. These cells were expanded for 1-2 passages and then sorted for CD45-, CD11b-, and F4/80-negativity to exclude contaminating macrophages. The sorted MEFs were split once after sorting to allow for recovery and used for experiments at p4-p7. Unsorted MEFs (p1-p5) were used in experiments where noted.

### Cytokines and chemicals

Growth factors were used at the indicated concentration. If not specified, CSF-1 and PDGF-BB were used at 50 ng/ml, TGF-*β* was used at 10 ng/ml, and WNT3A was used at 50 ng/ml. Chemical inhibitors were titrated based on published literature and used at the following concentrations: Rho activator CN03 2 ug/ml, TGF-*β* pathway inhibitor LY364947 1 uM, actin filament inhibitor LatA 500 nM, Rock inhibitor Y27632 10 uM. For experiments with chemical inhibitors, cells were plated at different cell densities overnight, treated with inhibitors, and collected at the indicated time for analyses of RNA expression, cellular signaling, or immunofluorescent imaging. For cells treated with CN03, all conditions were plated in complete DMEM first, then changed to serum-free complete DMEM overnight before applying CN03.

### Flow cytometry for cell quantification

MEFs were harvested from tissue culture plates by incubation with 0.05% Trypsin + EDTA. BMDMs were harvested from non-tissue culture treated plates by incubation with 3mM EDTA in PBS. MEFs and BMDMs in co-culture were harvested first by incubation with 0.05% Trypsin + EDTA, and next by cell scraping to remove any remaining attached cells. All cells were washed and transferred to round-bottom 96-well plates for FACS staining. Fluorochrome-conjugated antibodies against CD45 (clone 30-F11), CD11b (M1/70), and F4/80 (BM8) were purchased from eBioscience/ThermoFisher. Dead cells were excluded using ThermoFisher LIVE/DEAD Fixable Aqua Dead Cell Stain and binding to Fc receptors was blocked using CD16/CD32 (clone 93, eBioscience/ThermoFisher). Absolute cell numbers were calculated using counting beads (123count eBeads, ThermoFisher), which were added following harvest, immediately prior to running samples on the flow cytometer (50 uL of beads were added to 150 uL sample). Absolute cell numbers were calculated according to manufacturer’s instructions, using the following equation: ((# of live single cells acquired * 0.05)/(# of beads acquired * 0.15)) * eBead concentration. To correct for cell loss during harvest, we applied a standard curve adjustment based on the original cell number plated and the bead-calculated cell number. All samples were acquired on a Becton Dickinson LSR II Flow cytometer and analyzed using FlowJo.

### Cell isolation from liver

Livers from sacrificed mice were prepared by mechanical disruption, followed by 30 minute treatment with 2 mg/ml Collagenase Type 4 (Worthington Biochemical) in PBS at 37°C with continuous shaking. Digested tissues were mashed through 70 μm filters, layered in a 33% and 66% Percoll gradient (Sigma), and centrifuged at 3000 rpm for 30 min without brake. Cells at the interface were collected and analyzed by flow cytometry. Fluorochrome-conjugated antibodies against CD45 (clone 30-F11), CD64 (clone X54-5/7.1), and MerTK (clone DS5MMER) were purchased from eBioscience/ThermoFisher. Dead cells were excluded using Zombie Yellow Fixable Viability Stain (BioLegend) and binding to Fc receptors was blocked using CD16/CD32 (clone 93, eBioscience/ThermoFisher). Cell numbers were calculated using counting beads (123count eBeads, ThermoFisher) as described above.

### Assay to quantify growth sensitivity to space availability and growth factor

We used Click-iT Plus EdU Flow Cytometry Assay Kit to quantify proliferation of MEFs or BMDMs at different conditions. Briefly, MEF or BMDM monocultures were plated in 6-well tissue culture plates 1 day prior to EdU incorporation assay. EdU (10 mM) was added to the cells for 2 hr and the cells were harvested as above. Cells were first stained with surface antibodies and then EdU incorporation was detected using the Click-iT Plus EdU Flow Cytometry Assay Kit according to the manufacturer’s instructions (ThermoFisher). Samples were then acquired on a Becton Dickinson LSR II Flow cytometer and analyzed using FlowJo. For analysis of sensitivity to space availability, cells were plated at 50,000 to 1,000,000 per well in 6-well plates, which covers the range that proliferation changes linearly with cell density. In each experiment, the percentage of EdU^+^ cells was plotted against cell density, and the slope of linear regression was normalized to represent change of proliferation per 100,000 cells (*Δ*Edu+/100K). For analysis of sensitivity to growth factors, we treated cells with 0 - 20 ng/ml growth factors overnight, before analyzing using the EdU assay kit described above. To confirm the observed results were not due to insufficient growth signals, particularly for fibroblasts, cells were treated with high concentrations of CSF-1 and PDGF-BB up to 200 ng/ml. The proliferative effect of recombinant growth factors saturated at less than 10 ng/ml for both macrophages or fibroblasts. To quantify the sensitivity to growth factor, we calculated the max growth difference between control (without recombinant growth factor) and growth factor stimulated conditions. Multiple experiments were averaged to infer the sensitivity of growth to space availability or growth factors, where each dot represents one experiment.

### Western blot analysis

Cells were washed 1X with ice cold PBS before harvesting. Protein was extracted in 1x SDS buffer (62.5 mM Tris-HCl, pH 6.8, 2% SDS, 25% glycerol, 0.01% bromophenol blue) on ice followed by heating at 95°C for 10 minutes. Multiple wells of low density cells were combined during collection to reach similar total cell numbers at each cell density. Lysates corresponding to equal numbers of cells were loaded into 4-20% TGX Precast gels (Biorad) and ran with Tris/glycine running buffer (Biorad). Protein was transferred onto activated PVDF membrane (Millipore) using trans-blot Turbo system (Biorad), and then blocked using 5% BSA in TBST (20 mM Tris, 150 mM NaCl, 0.05% Tween 20) for at least 1 hour at room temperature. Primary antibody was incubated in 5% BSA in TBST at 4°C overnight at recommended dilution (anti-SMAD2/3 D7G7 1:1000, anti-phospho-SMAD2/3 D27F4 1:1000, anti-GAPDH 1:2000), and secondary antibody was incubated in TBST at room temperature for 30 min at 1:5000. Samples were washed 3x with TBST following each round of antibody incubation. Protein was visualized using ECL western blotting substrate (ThermoScientific) and autoradiography film. Densitometry was calculated using ImageJ software. Band intensity of interest was normalized to GAPDH for each sample, and averaged across biological replicates.

### Viral transduction

Expression vectors pMigR1-Cre/IRES-GFP (Cre-GFP) or pMigR1-IRES-GFP (GFP) were co-transfected into 293T cells with retrovirus packaging vector pCL-Eco using the Lipofectamine 2000 kit (Thermo Fisher Scientific, #11668019) according to the manufacturer’s instructions. After 24 hours, cell culture medium was changed to complete DMEM. After 24 hours, viral supernatant was collected, filtered through 70 μm cell filters and directly applied to low-passage unsorted MEFs, at 1:1 ratio to the existing medium. Successfully transduced MEFs were sorted based on GFP-positivity and allowed to rest for at least one passage before performing co-culture experiments.

### Immunofluorescence imaging

Cells were cultured in 8 well chamber slides and fixed in 4% paraformaldehyde and permeabilized using 0.1% saponin in blocking buffer (HBSS containing 3% BSA, 0.2% gelatin, and 0.02% NaN3) and stained with Rabbit anti-mouse YAP1 (Cell Signaling Technology, D8H1X), FITC-conjugated phalloidin, and Goat anti-rabbit IgG (H+L) Alexa fluor 594 secondary (ThermoFisher, A-11007). Hoechst 33342 was used to stain nuclei. Cells were mounted on microscope slides with ProLong Diamond Antifade Mountant (Molecular Probes). Imaging was performed with Leica AF6000 Modular System (fluorescent microscope) or Leica SP8 (confocal microscope). Confocal imaging was acquired using HC PL APO CS2 63x/1.40 OIL objective and the 405nm and Argon (488nm) lasers collected on different sequentials.

### Quantification of nuclear shape and actin fluorescence

Slides stained with phalloidin and DAPI were imaged with a Phalloidin corrected fluorescence intensity measured and calculated using Fiji (ImageJ) version 2.1.0/1.53c.(Schindelin et al., 2012) Cells were manually outlined using freehand selection around phalloidin staining. For each cell, area, mean gray value, and integrated density were collected for all cells and an area with no cells was collected for background signal quantification. Corrected fluorescence intensity calculated with the equation: Integrated density – (area of cell * mean gray value of background). Figures generated using ggplot2 in R with R Studio. Statistics calculated using Wilcox t test comparing all densities to the 10k density individually.

Nucleus quantification was obtained after collecting confocal z stacks with slices every 0.3um encompassing the entire nucleus of all cells fully within the field of view. 3D projections viewed using Imaris software (Bitplane), and surfaces were generated using DAPI fluorescence intensity. Statistics for all surfaces were exported and figures were generated using ggplot2 in R with R Studio. Length, width, and thickness were obtained using the object oriented measurements with bounding box length A, B, and C respectively. Statistics calculated in R using Wilcox t test comparing all densities to the 10k density individually.

### ECM and supernatant transfer

For ECM “transfer,” 200,000 unsorted MEFs were plated per well in 6 well plates in 10% FBS for 2 days. After 2 days, cells were removed using 25 mM EDTA in PBS and blasting by pipetting. Plates were then washed twice with PBS and fresh 20,000 or 200,000 MEFs were plated on top of decellularized matrix. Cells were harvested and RNA isolated after overnight culture. For supernatant transfer experiments, supernatant from either high density (200,000) or low density (25,000) MEF cultures was removed after 12 hours, spun down to remove any cellular debris, and added to MEFs plated at low density (25,000). Cells were harvested and RNA isolated after overnight culture.

### Gene silencing using siRNA

P1-P2 unsorted MEFs were lifted from plates using 0.05% Trypsin + EDTA. Each siRNA was incubated with OptiMEM while RNAiMax was incubated with OptiMEM for 5 mins at room temperature. The two solutions were mixed together dropwise and incubated for 20 mins. siRNA mixes were then plated in 12-well plates and 50,000 MEFs were added to each well. All siRNAs were used at a final concentration of 5nmol. Cells were harvested and RNA isolated after 3 days of incubation with siRNA.

### RNA isolation and qRT-PCR

RNA was purified from cells using Qiagen RNeasy columns with on-column DNAse digestion according to the manufacturer’s instructions. cDNA was reverse-transcribed with MMLV reverse transcriptase (Clontech) using oligo-dT20 primers. qRT-PCR was performed on a CFX96 Real-Time System (Bio-Rad) using PerfeCTa SYBR Green SuperMix (Quanta Biosciences). Relative expression units were calculated as transcript levels of target genes over 1/1000 of Actb. Primers used for qRT-PCR are listed in Table S1.

### RNA sequencing

Primary p0 MEFs were expanded for 1 passage and sorted to remove contaminating cells. Sorted MEFs were culture for 1 passage and plated at the following cell density 10,000/well (2,000/cm^2^), 50,000/well (5,000/cm^2^), 200,000/well (21,000/cm^2^), and 500,000/well (52,000/cm^2^) in DMEM. The highest cell density is close to the theoretical carrying capacity that we determined previously (Zhou et al., 2018). After overnight culture, adherent cells were washed twice with ice-cold PBS and collected with RLT buffer (Qiagen RNeasy kit). RNA was purified from cells using Qiagen RNeasy columns with on-column DNAse digestion according to the manufacturer’s instructions. Sequencing libraries were constructed following Illumina Tru-seq stranded mRNA protocol. Paired-end sequencing was performed with Next-seq 500 with either 76 or 38 bp from each end. Two sets of replicated densities were obtained from different batches of primary MEFs.

### RNAseq analysis

Illumina fastq files were downloaded from Illumina Basespace and were aligned with Kallisto program with default settings (Bray et al., 2016) against all cDNA transcripts in mouse genome annotation GRCm38 (ftp://ftp.ensembl.org/pub/release-90/fasta/mus_musculus/cdna/). The ENSEMBL IDs of each cDNA transcript were matched to the official gene symbols through BioaRt in R. The expression of each transcript is expressed in TPM (transcript per million). When multiple transcripts match to the same gene, the expression of the gene is calculated by summing the TPM of all matched transcripts.

### Differential expression analysis

Differential expression analysis was performed using sleuth program (Pimentel et al., 2017). This analysis returned 34475 genes in total in the mouse genome. Using an adjusted p-value of 0.05, we identified 3046 genes that are significantly differentially expressed between any two cell densities and have an expression higher than 2 TPM. To further identify genes that are differentially expressed biologically, we further selected genes that are activated or repressed more than 1.5-fold. Primary fibroblast cultures have significant biological variations between each preparation. To identify the genes consistently influenced by cell density, we performed the experiments with two separately isolated batches of MEFs. As expected, there are significant variations between biological replicates at baseline. We developed an automated correlation-based method to identify the genes that change with cell density consistently. To select a threshold for defining consistency between biological replicates, we computationally shuffled the data to generate a randomized background with no logical connection between each density for each gene. After ranking the genes from the least correlated to the best correlated, we calculated the accumulated gene counts as a function of the correlation coefficient. For the randomized dataset, this gives a linear line, because different degrees of correlation happen by chance (Figure S1C). For the experimental dataset, this gives a curve skewed towards high correlation coefficient (Figure S1C). We used the intersection between the randomized and experimental data curves as the threshold for selecting genes with consistent changes in expression. Intuitively, far more truly correlated genes are selected than genes with randomized expression at this correlation coefficient. Following these criteria, we identified 1950 genes as density-dependent genes. All fold changes are calculated using a pseudo count of 1 TPM to avoid dividing expression value by 0 and small expression values. These genes were then clustered using K-mean clustering based on log2(fold) changes in TPM expression. K-mean clustering was performed with customized codes using built-in functions in Matlab, until the clustering converged in 20 iterations. Clusters of 2 - 10 were screened and 7 was identified as the minimum number of clusters to characterize the general trend of expression changes at different cell densities.

### Sequence motif and function enrichment analysis

Clusters of genes were organized into different groups and analyzed with Homer using motifs.pl. For sequence motif enrichment, the analysis focused on gene promoter regions, defined by -1000 and +200bp from transcription start site, searching for DNA sequence motifs from 6 to 10 bp in length. For function enrichment analysis, annotated cellular signaling pathways were extracted from Kegg pathways and all significantly enriched terms (p <= 0.01) were curated based on enrichment statistics.

### Chromatin-immunoprecipitation and high-throughput sequencing

MEFs were crosslinked by adding formaldehyde to the medium to a final concentration of 1% in 15 cm plates with continuous, gentle rocking at RT for 10 minutes. The reaction was quenched by adding 125 mM final concentration of glycine, with continuous, gentle rocking at RT for 5 minutes. The cells were washed 3 times with ice cold PBS, scraped with 10 ml of PBS, and spun down for 5 minutes at 1,500 rpm at 4°C. For Chromatin-immunoprecipitation, for cell pellets of 10^6^ cells, 1 ml cell lysis buffer (10 mM Hepes pH 7.3, 85 mM KCl, 1 mM EDTA, 0.5% IGEPAL CA-630, 1x protease inhibitors (ThermoFisher Halt)) was used to resuspend the pellet and incubated on ice for 5 minutes. The lysate was centrifuged for 5 minutes at 4,000 rpm at 4°C, the supernatant was removed, and the pellet was resuspended in 0.3 ml of nuclear lysis buffer for sonication (10mM Tris pH 8.0, 0.5% N-lauroylsarcosine, 0.1% sodium deoxycholate, 100 mM NaCl, 1 mM EDTA, 0.5 mM EGTA, 1x protease inhibitors (ThermoFisher Halt)). Cell lysate was transferred to 1.5 ml Bioruptor Plus TPX microtube (Diagenode) and was sonicated for 4 rounds of 15 cycles of 30 seconds on/30 seconds off on high power (Diagenode Biorupter plus). After sonication, the chromatin was centrifuged for 15 minutes at max speed at 4°C and the supernatant was carefully transferred to a new tube and 1% Triton X-100 was added. 0.7 mL of ChIP dilution buffer (20 mM Tris pH 7.5, 0.5% Triton X-100, 100 mM NaCl, 1 mM EDTA, 1x protease inhibitor) was then added to the sonicated chromatin. 10% of the chromatin was taken as “Input”, and the remaining chromatin incubated overnight with anti-YAP1 antibody at 1:50 dilution (CST) overnight on a rotator at 4°C. Protein G Dynabeads (ThermoFisher 10004D) were washed 3 times with PBS + 0.5% BSA and then added to the chromatin and antibody mixture and rotated at 4°C for 3 hours. Protein G dynabeads were washed sequentially with low-salt wash buffer (20 mM Tris pH 7.5, 0.1% SDS, 1 % Triton X-100, 150 mM NaCl, 1 mM EDTA), and LiCl wash buffer (10 mM Tris pH 7.5, 1% sodium deoxycholate, 1% Triton X-100, 250 mM LiCl, 1 mM EDTA) 3 times, and washed once with TE to remove detergents and salts. Bound DNA was eluted twice by resuspending the beads with 125 ul of elution buffer (50 mM NaHCO3, 1% SDS) and incubated for 10 minutes at RT, followed by 3 minutes of vigorous shaking at 37°C. Eluted DNA was digested with proteinase K at 55°C for 2 hours and reverse-crosslinked at 65°C overnight. DNA was purified using the Qiagen MinElute PCR Purification Kit and ChIPseq libraries were generated using the NEBNext Ultra II DNA Library Prep Kit for Illumina. Sequencing was performed with Illumina NextSeq500 for paired-end 38bp reads.

### ChIP-seq analysis

Reads from Illumina paired end fastq files were mapped to the mouse genome (mm9) using Bowtie(Langmead et al., 2009) with the options “-S -q --best -m 1 -p 20 -v 1 -a -strata” to generate SAM files. Read duplicates were removed and BAM files were generated with the Picard toolkit (http://broadinstitute.github.io/picard.). Bigwig files were generated using Homer software(Heinz et al., 2010) using the option “-fsize 1e20” and uploaded to UCSC genome browser. Enriched chip regions were identified using MACS2(Zhang et al., 2008) using the options “’-f BAMPE --bw 200 -B -g mm”. Transcription factor motifs in the enriched chip regions were identified also using Homer software using the options “-mask -size 200 -mis 3 -S 30 -len 8,10,12”. Genes were associated with the enriched chip regions by locating the closest gene also using Homer software.

### Analysis of histone modifications

Histone modification (H3K4me1, H3K4me3, K3K27ac) data of mouse embryonic fibroblasts were obtained from Mouse Encode http://www.mouseencode.org/. H3K27ac data of human skeletal muscle fibroblasts (HSMM), endothelial cells (HUVEC), myeloid leukemia cells (K562), B-lymphoblastoid cells (GM12878), stem cells (hESC), epidermal keratinocytes (NHEK), and human lung fibroblasts (NHLF) were obtained from the human ENCODE project (https://www.encodeproject.org/). The tracks of HSMM, HUVEC and K562 were shown as examples.

### Modeling the effect of CSF1 production rate on macrophage numbers

To model the effect of varying the production rate of CSF1 in fibroblasts, we used mathematical models describing cell circuit communication between macrophages and fibroblasts developed in our previous work (Adler et al., 2018; Zhou et al., 2018). Model parameters are as described before except the rate of CSF1 production. We numerically solved the ODE system to extract the steady-state level of macrophage and fibroblast numbers at different CSF1 production rates. This is performed with the NDSolveValue and FindRoot functions in Mathematica 12.1.1.0.

### Single cell analysis

We analyzed single-cell RNA-seq data of human lung fibroblasts from Adams et al..(Adams et al., 2020) The data includes 1051, 786 and 4329 cells from control, COPD, and IPF patients, respectively. To preprocess the data, we used ‘Sanity’ – a recently developed method to normalize single-cell data and to infer the transcriptional activity of genes. Sanity is a unique Bayesian procedure for normalizing single-cell RNA-seq data from first principles.(Breda et al., 2019) Following the Sanity normalization, for each comparison between datasets, we removed genes with low expression (log10(average expression) > -11) and low expression variance (standard deviation > 0.05) across single cells. In total, 6934 genes in control fibroblasts, 7011 genes in integrated control and COPD fibroblasts, and 7376 genes in integrated control and IPF fibroblasts passed the selection thresholds. For the control fibroblasts, we observed a fraction of cells with extremely low gene expression (log10(average expression) < -9.9). After removing these outlier cells, the control samples included 945 cells for downstream analysis.

After preprocessing the data, we first established landmark genes whose expression depends on density of cells and shows significant variability in the single cell dataset. We considered z-scored expression in the single cell datasets for the top density dependent genes that show more than 2-fold differential expression between low (10K) and high (500K) cell densities. We identified 17 landmark genes for the control samples, including *Serpine1*, which is representative of genes expressed at a higher level at low cell density, and DCN, representative of genes expressed at a higher level at high cell density. Since the relative expression of these landmark genes correlates with the density of cells, we then defined a density score for each cell as the difference between the average expression of the high-density genes and the average expression of the low-density genes. For example, if a cell expresses high-density genes at a higher level than low-density genes, it will be given a high score for high density, and vice versa. We next sorted the cells according to their density scores and inferred the genes that are highly variable across this density axis. We repeated this analysis for the control fibroblasts alone, the control and COPD fibroblasts, and the control and IPF fibroblasts.

### Agent-based Modeling of the MPs FBs circuit

The agent-based modeling simulation was built in NetLogo, a programming language and IDE developed by Uri Wilensky and Northwestern University. In these simulations, we utilize the Patch type, which represents an individual square on the grid of the simulation space, entirely made of Patches. Blue patches emit a red signal (*R*, promoting red proliferation) and a self-promoting blue signal (*B2*). Red patches emit a blue signal (*B1*, promoting blue proliferation). At each timestep, the patches calculate the value of the signal perceived in their neighborhood. A patch’s neighborhood is defined as a circular radius centered around the patch itself and is a parameter that can be varied. The simulations presented in Figure 5 were run with a neighborhood radius of five patches. The calculation of signals for each patch at each timestep is given by the following equations.

With density-dependent regulation of red signal: 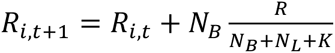, where *R_i,t_*_+1_ is the amount of red signal perceived by patch *i* at timestep t, *N_B_* and *N_L_* are the number of blue and lime patches in the neighborhood of patch *i*, *R* is the amount of red signal produced per patch, and *K* is a parameter that controls the effect of the density-dependence of blue cells on the red signal. Without density-dependent regulation of red signal: *R_i,t_*_+1_ = *R_i,t_* + *N_B_R* . Since blue signal is not subjected to density-dependent regulation, the amount of blue signal perceived by the patches at each timestep is the same in both cases: *B_i,t_*_+1_= *B_i,t_* + *N_R_B*_1_ + *N_B_B*_2_.

Depending on the value of signal, the patch decides to divide, stay the same, or die. In case of division, the patch randomly selects an empty (white) neighboring patch and changes it to be the same color. In the case of cell death, the patch turns empty (white), allowing for other neighboring cells to divide into its space.

### Data availability

RNA sequencing data of fibroblasts at different cell densities and at various stress conditions from this study will be deposited to the GEO repository and are accessible through accession number GSEXXXX and GSEXXXX. ChIPseq data of endogenous YAP1 MEFs have been deposited to GSE184774. Single cell RNAseq data of human lung fibroblasts are GSE136831. ChIPseq data of SMAD3 in MEFs were obtained from GSE85177. H3K27ac, H3K4me, H3K4me3 of MEFs were obtained from GSM1000139, GSM769028 and GSM769029 respectively. Human H2K27ac of fibroblasts, endothelial cells, and myeloid leukemia were obtained from GSM733755, GSM733691, and GSM733656. Customized scripts for density-dependent analysis, clustering in Matlab, imaging quantification in R, and circuit modeling in Mathematica are available upon reasonable request.

## Supplemental Information

**Figure S1.**
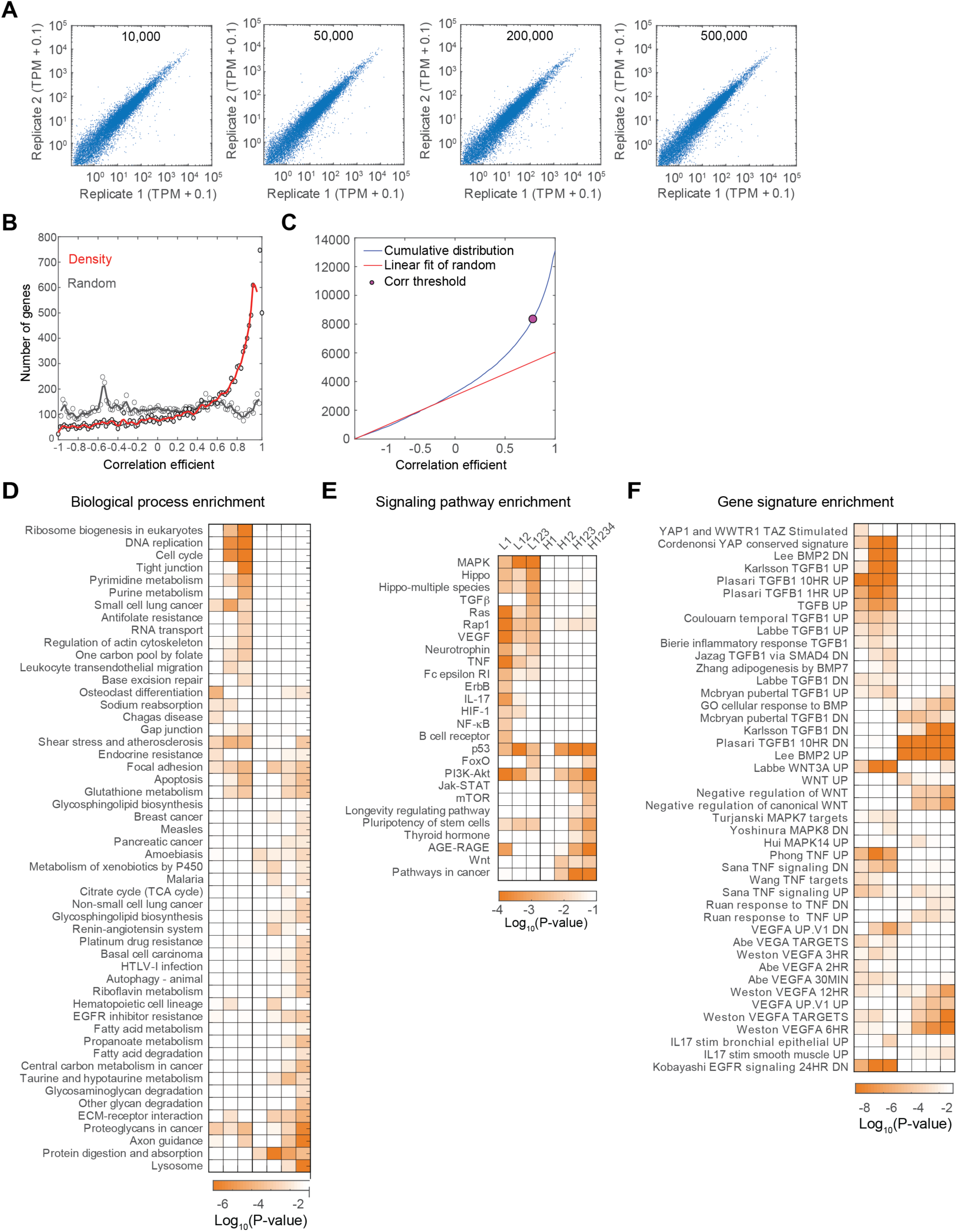
Transcriptional analyses of density-dependent expression programs. A. Transcriptome correlation of biological replicates at the indicated cell density. 0.1 TPM is added to each gene. B. Distribution of gene-specific correlation coefficients between two sets of randomized RNAseq data or biological replicates at matched cell densities. Expression data were generated randomly using a bootstrap method by shuffling genes and samples. Correlation coefficients of all genes are binned every 0.02 unit, and the sliding-window average is shown as colored curve. C. Cumulative distribution of the coefficients between two sets of biological replicates. Linear fit was generated using the distribution of coefficients that are less than 0. Linear fit of the cumulative distribution of correlation coefficients calculated using randomized data set is shown in red. The pink circle marks the threshold for consistent density regulation. D. Enrichment of biological processes in groups of genes induced at low or high density. E. Enrichment of signaling pathways in groups of genes induced at low or high density. F. Enrichment of molecular signature in groups of genes induced at low or high density.

**Figure S2.**
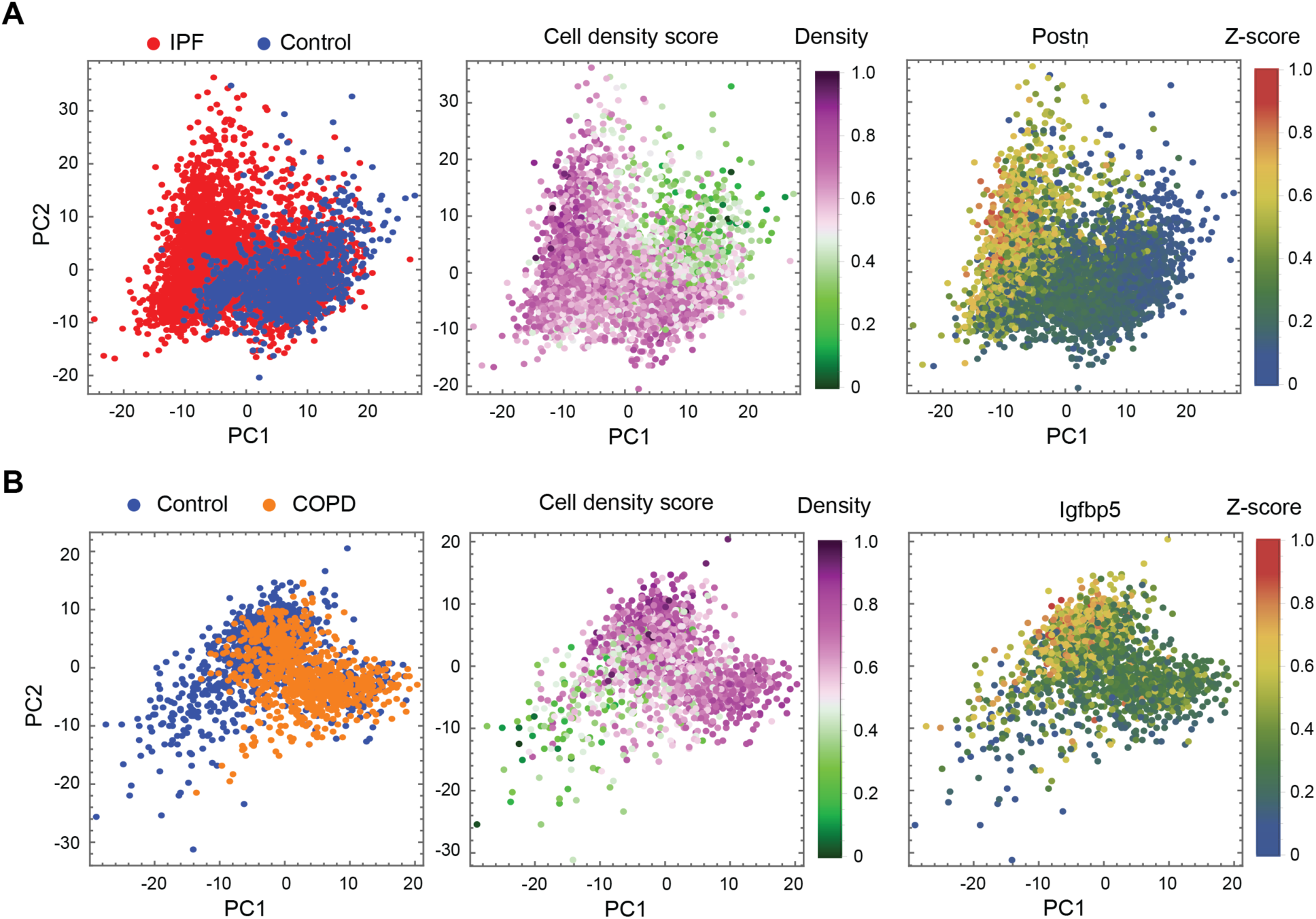
Single-cell reconstruction based on landmark genes reveals new densitydependent genes. A. Single-cell RNA-seq (scRNA-seq) data of fibroblasts from control (blue) and IPF (red) lungs projected on the first two principal components (left panel). Cells are colored by their density score, defined by the average expression of landmark density dependent genes (middle panel). POSTN is highly expressed in cells with high density scores (right panel). B. Same as A but with fibroblasts from COPD lungs in orange (left panel), and expression of IGFP5 (right panel).

**Figure S3.**
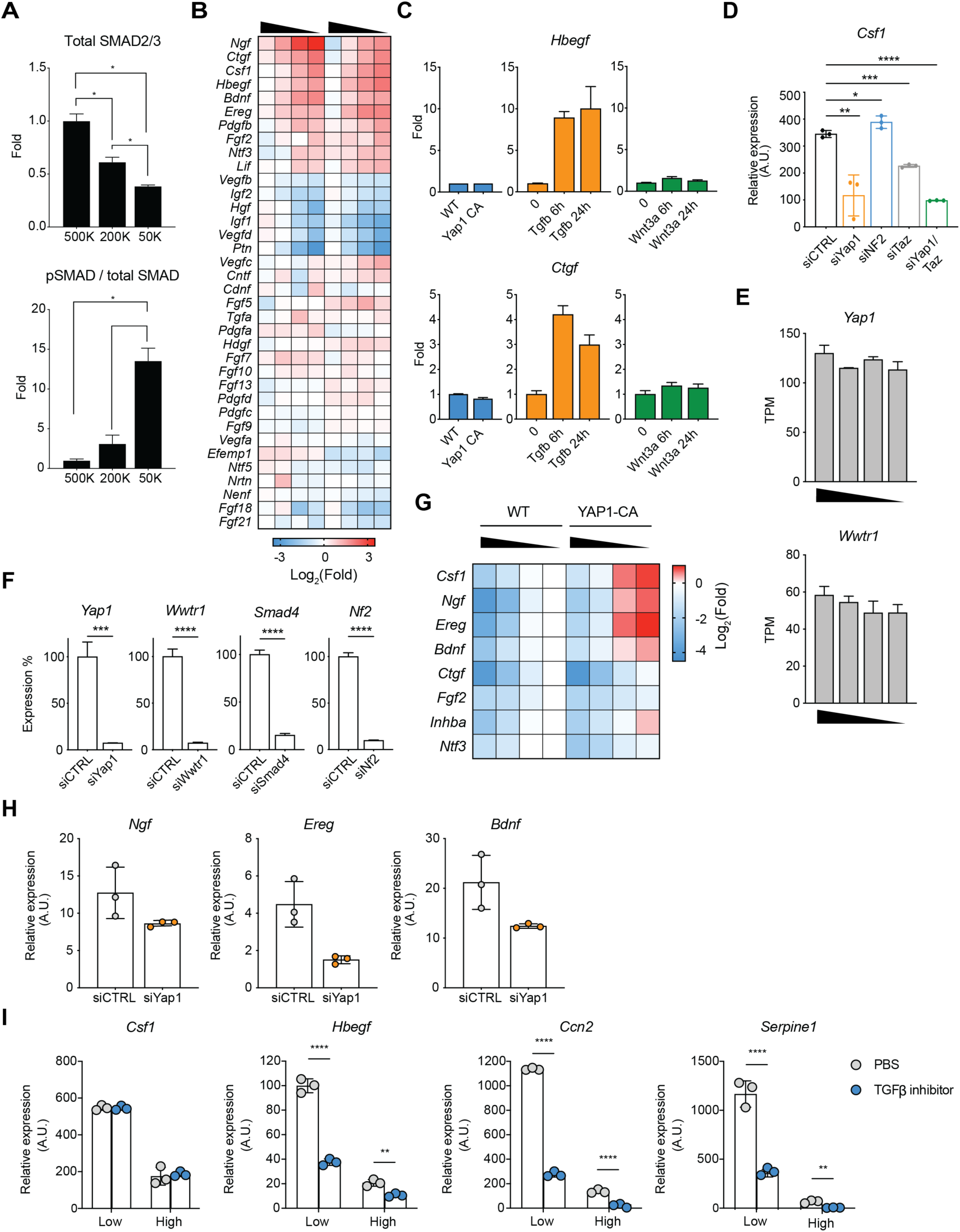
YAP1-dependent control of growth factor expression. A. Quantification of total Smad2/3 proteins, and the ratio between phosphorylated and unphosphorylated SMAD proteins at different MEF densities as measured by Western blot. Cells were analyzed after overnight culture at the specified cell densities. B. Heatmap showing the expression of growth factors in MEFs at different densities. Growth factors with TPM > 5 are shown. C. *Hbegf and Ctgf* expression in Yap^CA^ MEFs plated overnight, or MEFs treated with recombinant Tgfb or Wnt3a for 6 or 24 hours. D. *Csf1* expression in MEFs 3 days after transduction with siRNAs targeting *Yap1*, *NF2*, or *Wwtr1* (Taz) alone, *Yap1* and *Wwtr1*z together, or scrambled control siRNA (siCTRL). E. *Yap1* and *Wwtr1* expression in MEFs plated at decreasing cell densities, calculated as TPM from RNAseq data. F. siRNA knock-down of *Yap1*, *Ta*z, *NF2* and *Smad4* 3 days after siRNA transduction. G. Heatmap showing RT-qPCR quantified expression of selected growth factors at different cell densities in control and YapCA MEFs. H. RNA expression of *Ngf*, *Ereg*, and *Bdnf* 3 days after transduction with siRNAs targeting *Yap1*. I. RNA expression in MEFs cultured at low or high density after 4 hours treatment with 1uM TGF-β inhibitor, LY364947.

**Figure S4.**
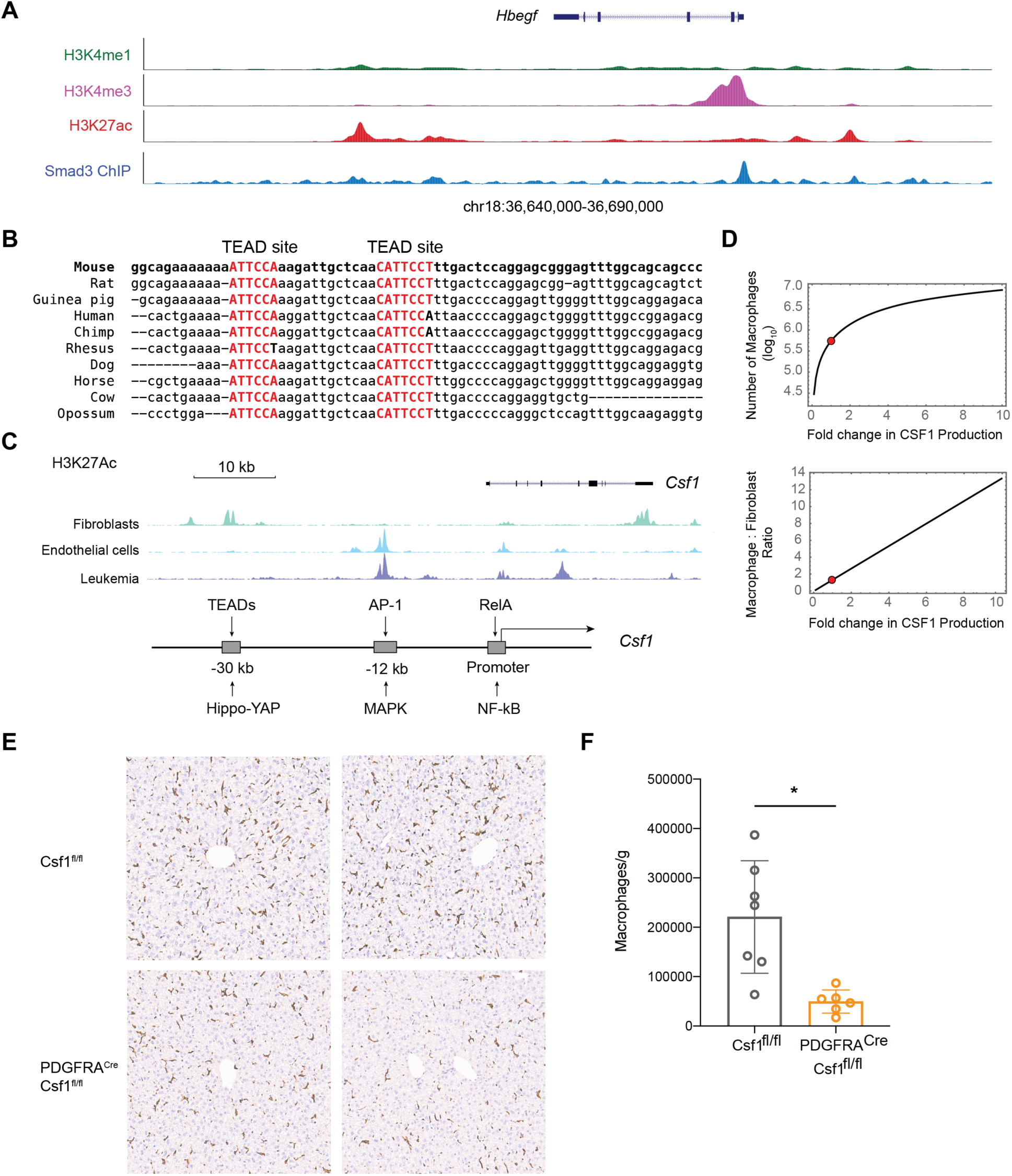
Regulatory elements at the *Csf1* and *Hbegf* loci and control of macrophage numbers by stromal cells. A. Genomic tracks displaying ChIP-seq of histone modifications (H3K27ac, H3K4me1, and H3K4me3) and Smad3 binding at *Hbegf* gene locus in MEFs. B. Alignment of genomic sequences within *Csf1* enhancer showing conservation of TEAD binding sequences among mammalian species. C. ChIP-seq tracks of H3K27ac in human cells (top) and illustration of enhancers and promoter at *Csf1* locus. H3K27ac data were obtained from the human ENCODE project. Diagram illustrates the known signals that regulate Csf1 expression (bottom). D. Cell circuit modeling prediction of macrophage numbers and macrophage to fibroblast ratios as a function of CSF1 production rate. The red circles mark the data measured using wild type cells. E. Representative immunofluorescence images of F4/80 staining (brown) in liver sections of Csf1^fl/fl^ and Pdgfra^Cre^Csf1^fl/fl^ mice. F. Number of liver macrophages in Csf1^fl/fl^ and Pdgfra^Cre^Csf1^fl/fl^ mice per gram of liver tissue. Cells are gated on live CD45^+^CD64^+^MerTK^+^ cells.

**Figure S5.**
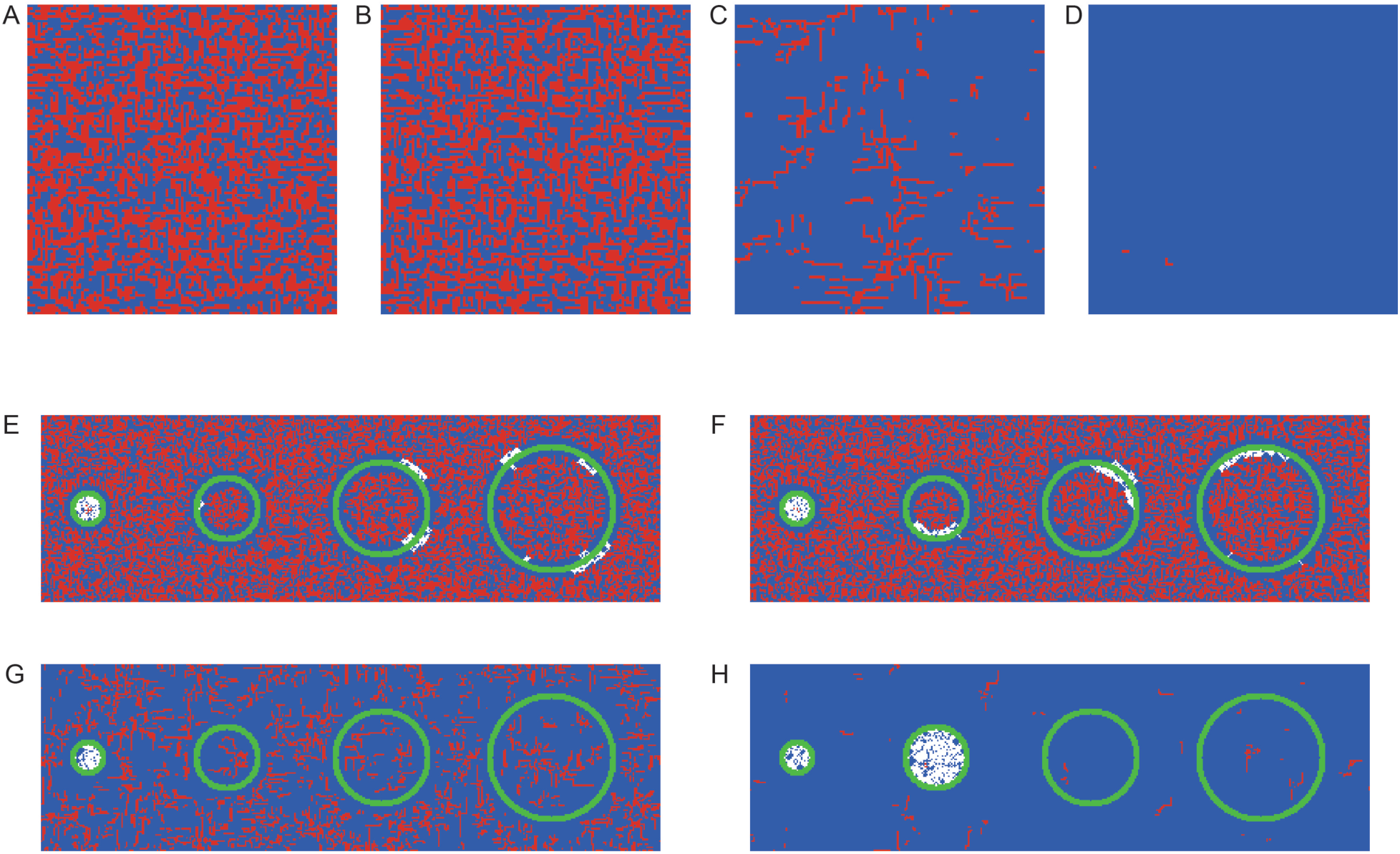
Effect of parameters on agent-based models. A-D. Equilibrium arrangements of red and blue cells following simulations that start with 17.5% red and 17.5% blue cells that are randomly placed on the 2D grid. The simulations are with a different value for the parameter K that controls the effect of density-dependent regulation of the red signal: K=1 (A), K=3 (B), K=5 (C) and K=7 (D). As the density-dependent regulation of red signal is increased, red cells cannot proliferate leading to their depletion and to the takeover of blue cells which are able to proliferate due to their autocrine blue signal (D). E-H. Equilibrium arrangements of red, blue, and inert (lime) cells following simulations that start with 20% red and 20% blue cells that are randomly placed on the 2D grid, and with K=1 (E), K=3 (F), K=5 (G) and K=7 (H).

**Figure S6.**
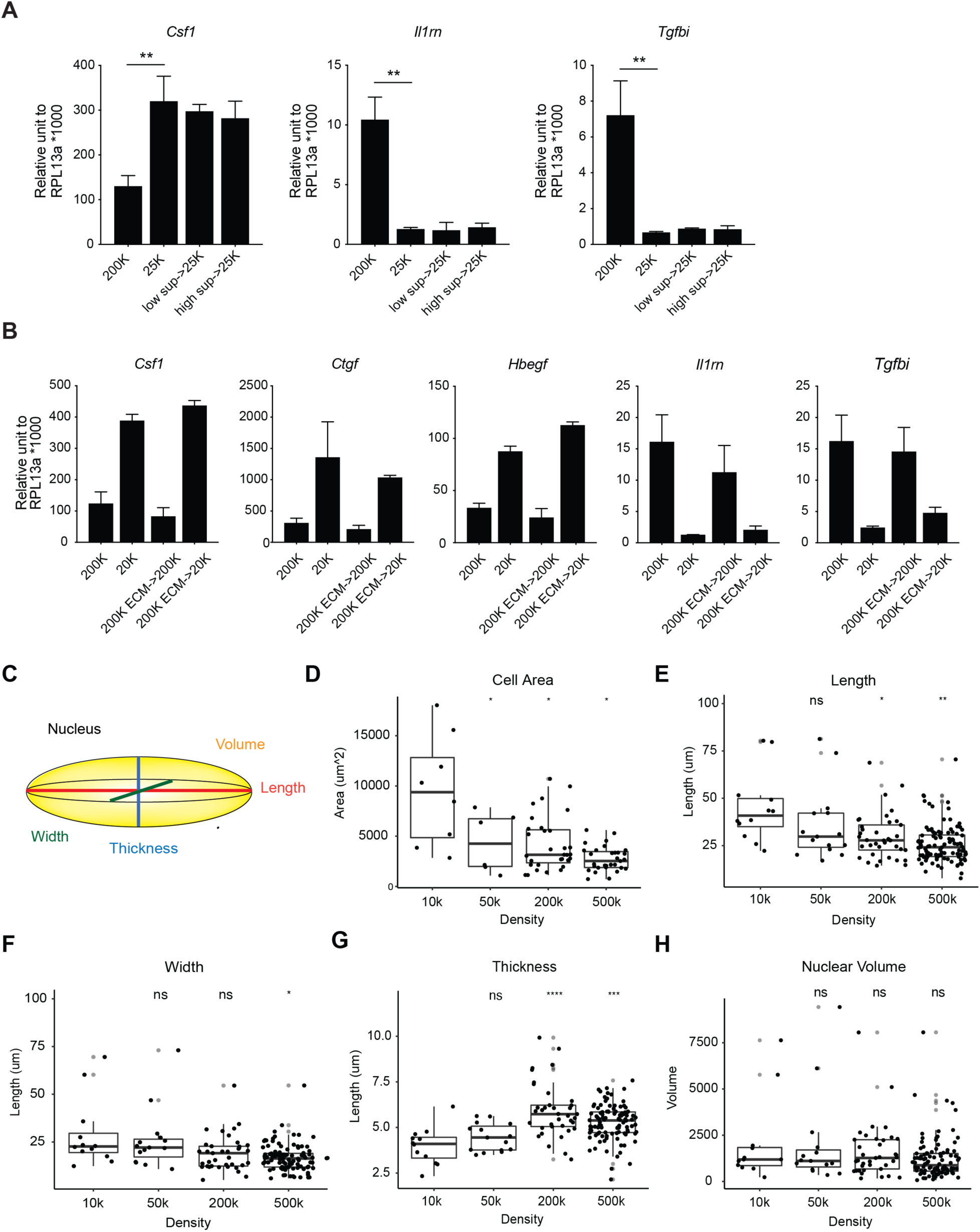
Cellular mechanisms controlling density-dependent expression programs. A. Expression of growth factors in MEFs plated at low or high density and cultured overnight in conditioned supernatant. Two-tailed t-test was applied to all statistical analyses, *p<0.05, **p<0.01. B. Expression of growth factors in MEFs plated on decellularized extracellular matrix from low or high density cell cultures. C. Graphical representation of the measurements for size and shape of nuclei. D-H. Quantification of cross-sectional cell area (**D**), length (**E**), width (**F**), thickness (**G**) and volume (**H**) of nuclei. Nuclear length is measured as the longest axis of the nucleus and nuclear width is quantified along the axis perpendicular to nuclear length in the *xy* plane. Nuclear thickness is defined as the shortest axis of the nucleus, typically along the z axis. Each point represents one nucleus. Statistical significance is performed with Wilcox t test. ns p>0.05, *p<0.05, ** p<0.01, *** p<0.001, **** p<0.0001.

**Figure S7.**
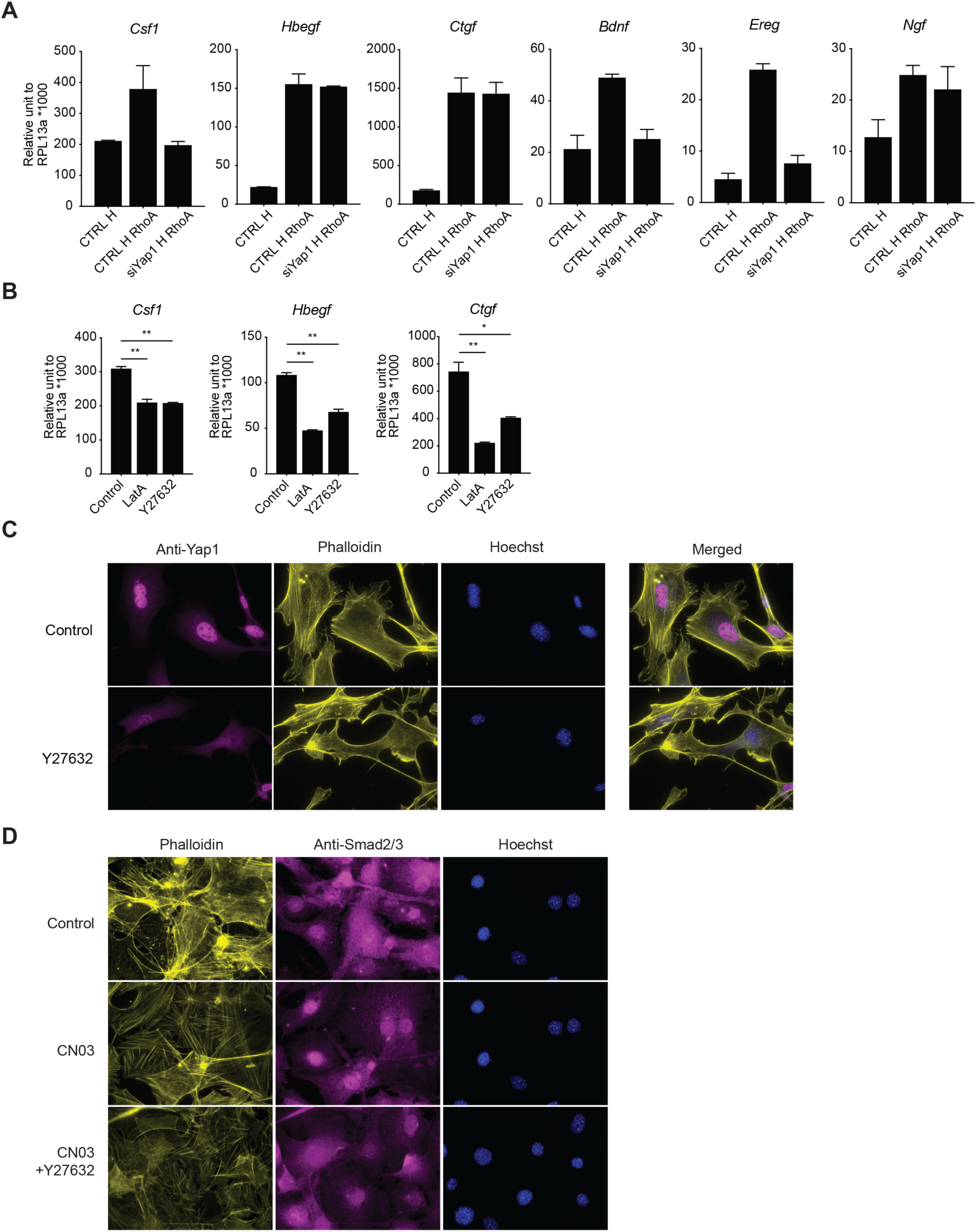
Actin-dependent mechanisms regulate activation of YAP1 and SMADs. A. Growth factor expression in MEFs plated at high cell density after treatment with RhoA activator (CN03 peptide) for 4 hours with or without 3 day transduction with siRNA targeting *Yap1* prior to peptide treatment. B. Expression of growth factors in MEFs treated with Latrunculin A (LatA) or ROCK inhibitor (Y27632) for 4 hours. C. Localization of YAP1 in MEFs plated at low density (equivalent to 50,000/well) and treated with Y27632 for 4 hours. D. Localization of SMAD proteins (total Smad2/3) in MEFs plated at high cell density (equivalent to 500,000/well) and treated with RhoA activator (CN03 peptide) or RhoA activator in combination with ROCK inhibitor for 4 hours.

## Notes

### Competing Interest Statement

The authors have declared no competing interest.

